# A foundation model enables prediction of natural product molecular properties, bioactivity, and structural similarity from biosynthetic gene cluster sequence

**DOI:** 10.64898/2026.07.05.736569

**Authors:** Allison S. Walker

## Abstract

Genome mining is a powerful technique in natural product discovery, where biosynthetic gene clusters that are likely to produce novel or desirable natural products are identified through bioinformatic analysis. There are many more predicted biosynthetic gene clusters than can easily be experimentally characterized. Additional computational methods to prioritize biosynthetic gene clusters by the bioactivity, structural properties, or novelty of the product would make genome mining more efficient. Multiple machine learning/artificial intelligence models have been developed to predict product properties from biosynthetic gene cluster sequence, but they are limited by small quantities of training data. Model pretraining with unlabeled data is a powerful technique to develop models that can learn on a limited amount of labeled training data. Biosynthetic gene clusters are well suited to this strategy because there are many predicted clusters with only a small percentage being characterized. This paper reports BGC-MLM, a foundation model that is pretrained with a masked language task on predicted biosynthetic gene clusters and then fine-tuned for downstream applications including prediction of product structural class, bioactivity, chemical properties, counts of functional groups, and chemical fingerprint. Comparison to a model trained without pretraining shows that pretraining generally improves performance. BGC-MLM shows better or similar performance to existing specialized methods for these tasks, demonstrating its utility as a foundation model for natural product genome mining.

## Introduction

Natural products are historically an excellent source of therapeutic molecules with complex structures that serve as an inspiration for chemists.^1^ Natural product genome mining, or the process of searching genomes for biosynthetic gene clusters (BGCs) that encode the enzymes necessary for natural product biosynthesis, has revealed that we have only just tapped the surface of natural product diversity.^2,3^ It is costly and time consuming to isolate the product of a specific BGC – this often requires either heterologous expression of the BGC, genetic manipulation of the target strain, or screens of growth conditions or small molecule elicitors.^4^ Given the large number of putative BGCs, it is not feasible to determine the structure and function of the products of all BGCs. Therefore, methods for prioritizing BGCs based on the novelty, structural features, or bioactivities of their products are needed. Various artificial intelligence/machine learning (AI/ML) methods have been reported for predicting bioactivity,^5–7^ structural class,^8–11^ or even the full structure of a natural product from its BGC.^12^ However, these methods, in particular bioactivity and full structure prediction, are limited by low accuracy due to small training sets.^5,6^ In addition, PRISM is limited by its reliance on a predetermined list of known tailoring reactions,^12^ preventing it from correctly predicting products for BGCs that may harbor novel tailoring enzymes. Given the wealth of unlabeled BGC data, models that leverage pretraining can likely learn patterns in BGCs to improve accuracy for downstream applications.

Masked language models such as bidirectional encoder representations from transformers (BERT) first emerged to process natural language.^13^ These methods were then adapted for scientific applications including protein and genome language models.^14,15^ Masked language models are pretrained by masking tokens and training a neural network to predict the masked tokens based on the other tokens in the sequence. This allows the model to learn patterns in the pretraining sequences. The model can then be fine-tuned to predict properties of the sequence such as sentiment for natural language^13^ or protein function for protein language models.^16^ If the same pretrained model can be repurposed for multiple applications, they are referred to as a foundation model.^17^ This is advantageous in applications where unlabeled data is abundant but labeled data is rare. BGCs and natural products are an excellent example of such an application. There is a wealth of sequencing data and putative BGCs can be easily identified by genome mining software such as antiSMASH.^18^ But the products of the vast majority of BGCs are unknown.^2^ In addition, bioactivity is unknown even in some cases where both the BGC and product are known.^5^ This lack of data makes it challenging to train accurate machine learning models that predict natural product structure, molecular properties, and bioactivity. Pretrained masked language foundation models can likely address this challenge.

There have been several masked language or related models described for BGCs. BiGCARP uses masked language model pretraining to develop a model that was later fine-tuned to distinguish BGCs from non-BGCs and predict product class, however it is only trained on 127,000 BGCs, which is relatively small on the scale of total predicted BGCs.^11^ Another method, BGC-Chemical Co-Embedding (BCCoE), was built on BiGCARP embeddings to link BGCs and products by developing a BGC and molecule co-embedding space.^19^ Another model also used masked language pretraining, based on RoBERTa, predicts product class and can suggest additional domains to add to BGCs for BGC design, but was pretrained on either only 239,021 BGCs, or larger genomic datasets that were not limited to BGCs.^20^ DeepSeMS uses a transformer architecture to predict products from BGCs but its training set is limited to an augmented set of characterized BGCs.^21^ BGC-Prophet works on ESM2 embeddings at the gene, rather than domain, level to identify and predict product classes of BGCs but is trained on an augmented BGC dataset based only on characterized BGCs, along with more negative examples.^21^ BGC-MAC and BGC-MAP predict product class and link BGC-and structure, respectively, which both rely on ESM2 to embed protein domain sequences.^22^ These models rely only on characterized BGCs and there is no pretraining, beyond that which is used to create ESM2, which only applies to protein or protein domain sequences and not entire BGCs. Similarly, BGCat predicts product class but does not have pretraining, beyond the pretraining done to create ESM-C which is used to create gene embeddings.^10^ Therefore, a model that leverages pretraining on many uncharacterized BGCs, but not other types of genomic sequences, and that is applied to many downstream tasks has still yet to be investigated.

This work reports BGC-MLM, a pretrained foundation model for BGCs. BGC-MLM is pretrained on almost 900,000 predicted bacterial BGCs and shows good performance at predicting masked tokens, demonstrating that it learns the patterns of BGCs. BGC-MLM was fine-tuned for a variety of downstream tasks including prediction of product structural class, bioactivity, molecular properties, number of various functional groups present, and prediction of the product’s molecular fingerprint. The predicted fingerprint can be used to match BGCs to their product and vice versa, comparing the similarity of BGCs, and clustering them. For most tasks, pretraining improves performance compared to a model without pretraining demonstrating the utility of this approach. BGC-MLM performed better than or on par with existing specialized methods, with the exception of BGC clustering where clustering of BGCs by their BGC-MLM predicted fingerprint did not perform as well as state-of-the-art BGC clustering methods.

## Results and Discussion

### Model Pretraining

A masked language model (BGC-MLM) was pretrained on a set of 897,008 bacterial BGCs not on the contig edge predicted by antiSMASH version 5,^23^ a commonly used genome mining software. While newer versions of antiSMASH are available, antiSMASH version 5 was used because our previously constructed dataset of BGCs were available for pretraining,^24^ and rerunning predictions with a newer version of antiSMASH would be computationally expensive for minimal expected gain.

BGCs are represented as a sequence of PFAMs, with additional tokens to indicate the start and end of the BGC and to pad the sequence to make all BGCs the same length. The masked language model was constructed with an architecture and training procedure based on BERT (Figure 1).^13^ A test set of 164,125 BGCs with fewer than 70% of PFAMs shared with any single training BGC was used to assess the ability of BGC-MLM to accurately predict a masked PFAM. Hyperparameters including the dimensions of the hidden representation, the number of layers and transformer heads, and dropout rate were varied and models were assessed based on their average accuracy and top-10 accuracy on both the training and test set. Changes in hyperparameters had very little impact on accuracy, with some improving by some metrics while getting worse at others (Table S1). Without a clear best model, two different models were chosen for subsequent fine tuning, which will be referred to as “small” (25,245,220 parameters) and “large” (102,936,100 parameters) that differ based on the size of their hidden representations and number of attention heads (768 vs 1792 for the hidden dimension and 12 vs 14 for the number of attention heads). Four additional “small” and “large” models with identical hyperparameters but different random seeds were trained.

**Figure 1.**
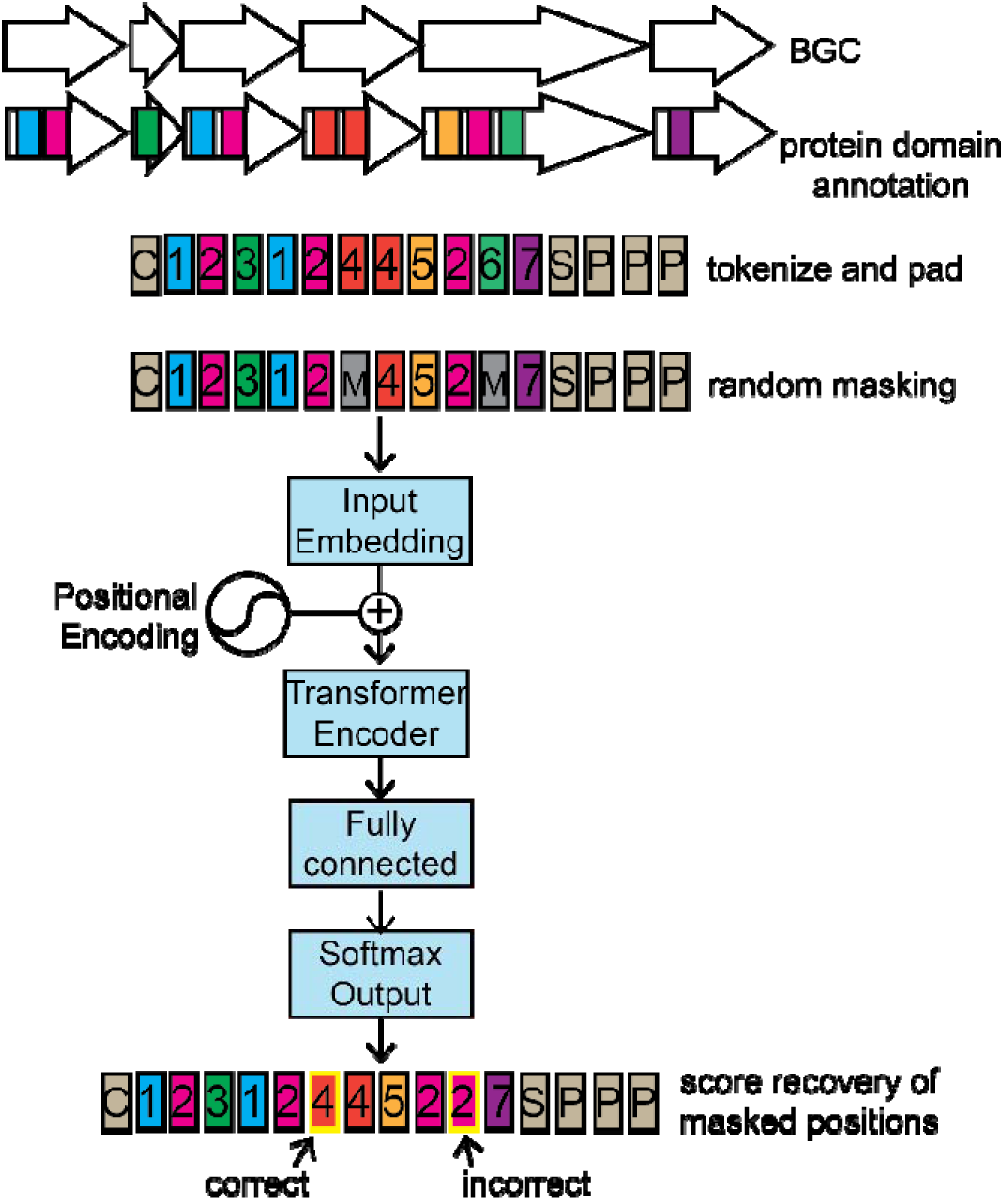
Illustration of BGC-MLM architecture. BGCs are represented as sequences of PFAM domain annotations and models are pretrained on a masked language task.

### Prediction of Natural Product Class

The ability of BGC-MLM models to predict natural product class was tested. The MIBiG database contains broad classifications of biosynthetic classes.^25^ Because a single BGC can belong to multiple biosynthetic classes, predicting the MIBiG label was treated as a multi-label classification problem. The possible classes are polyketide synthase (PKS), nonribosomal peptide synthetase (NRPS), RiPP (ribosomal), terpene, saccharide, and other. To construct the multi-label classifier, the BGC-MLM model was used to embed the sequence by mean pooling of the final layer and passed to a single fully connected layer and then an output layer. The MIBiG dataset was split with a stratified split to ensure the same distribution of classes in the training and test sets. Models were trained in three ways, one using frozen pretraining weights, where only the two layers added to construct the classifier are fit to the data (“frozen”), another where pretraining weights are used as a starting point but not frozen (“finetune”), and a final one where the model was trained from scratch not using pretrained weights (“scratch”).

All models performed well on the test data for this task with areas under the receiver operator characteristic curve (AUROC) ranging from 0.91 to >0.99 and areas under the precision recall curve (AUPRC) ranging from 0.72 to >0.99 (Figures 2, S1, Table S3). Performance varied by class, especially for the AUPRC metric, with NRPS, PKS, ribosomal peptides and terpenes being classified relatively well (AUPRC 0.81 to >0.99) and saccharide and other performing worse (AUPRC 0.72 to 0.88). The small, fine-tuned model performed the best by both AUROC and AUPRC for the plurality of classes (5 and 3 classes, respectively), although improvements over other models were generally small enough that they are likely not meaningful.

**Figure 2.**
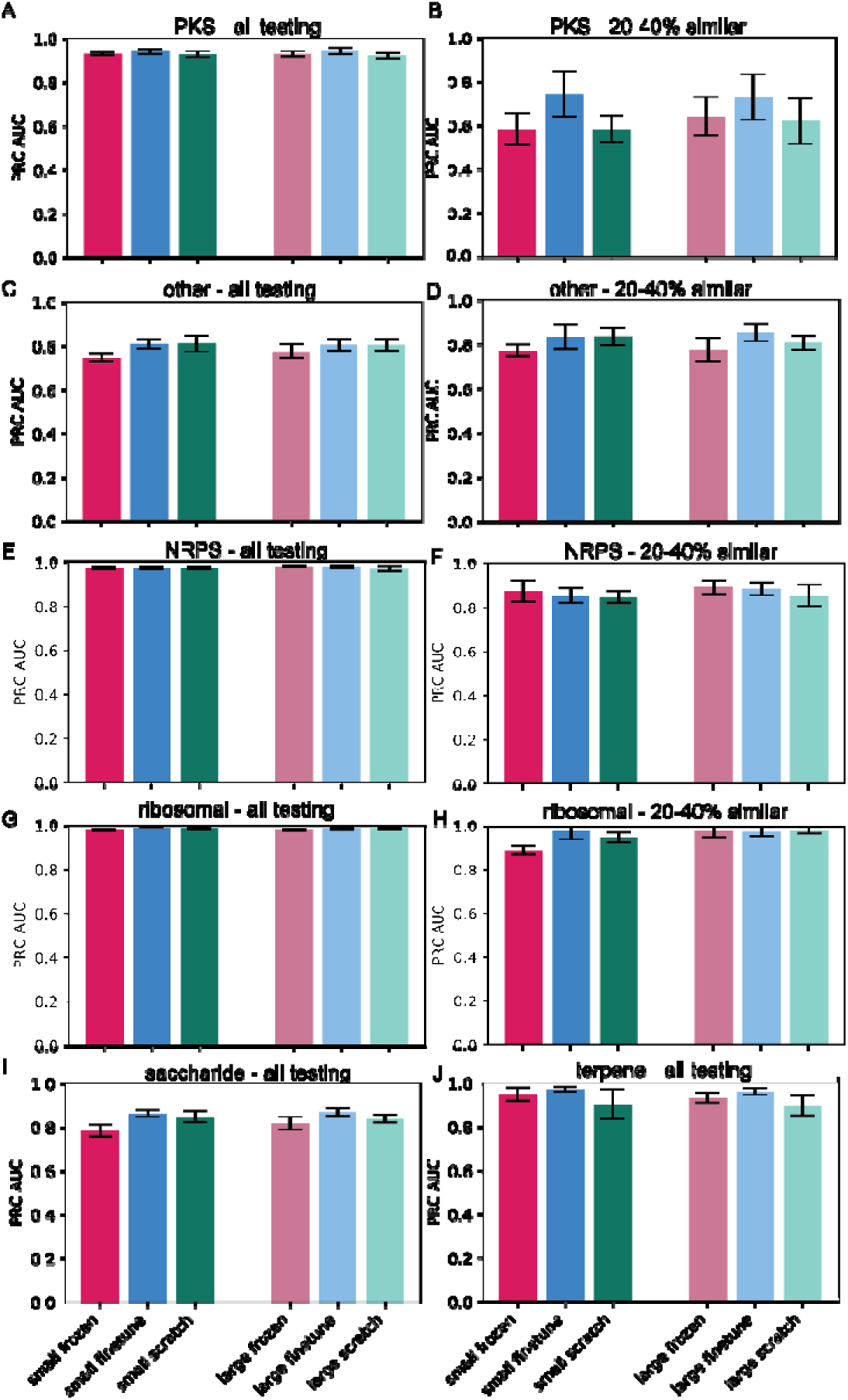
AUPRC by class for MIBiG classification task. **A)** PKS classification for entire test dataset **B)** PKS classification for BGCs with 20-40% similarity shared with finetuning set **C)** other classification for entire test set **D)** other classification for BGCs with 20-40% similarity shared with finetuning set **E)** NRPS classification for entire test dataset **F)** NRPS classification for BGCs with 20-40% similarity shared with finetuning set **G)** ribosomal peptid classification for entire test data set **H)** ribosomal peptide classification for BGCs with 20-40% similarity shared with finetuning set **I)** saccharide classification for entire test dataset **J)** terpene classification for entire test dataset. Ther were insufficient BGCs with 20-40% similarity to the finetuning set to calculate metrics for saccharides and terpenes for this set. Error bars indicate standard deviation between models trained with different random seeds.

To assess these models for overfitting, the test set was further divided by the percentage of PFAMs shared with the fine-tuning BGC in 20% increments. There were not enough BGCs of different classes with <20% shared PFAMs in either the pretraining or finetuning splits to assess performance. For BGCs in the 20-40% shared PFAMs with the fine-tuning set, performance decreased for some classes but remained better than what would be expected for a random model. NRPS, ribosomal peptides, and terpenes retained high performance (>0.8) (Figures 2, S2, Table S4). There was no consensus best model, but for the PKS classifier which showed the largest decrease in performance by AUPRC relative to the full test set, the fine-tuned models showed notably stronger performance than the frozen and scratch models. This supports the hypothesis that pretraining followed by fine-tuning allows models to better generalize across BGCs. For BGCs with 20-40% of PFAMs shared with the most similar pretraining example, performance was generally higher than for the BGCs with 20-40% of PFAMs shared with a BGC from the fine-tuning set (worst performing model had AUPRC of 0.56 compared to 0.47) (Figures S3-4).

Recently, two AI models for predicting more fine-grained molecular classes from BGCs have been reported, BGCat,^10^ which classifies NPClassifier classes,^26^ and CHAMOIS,^9^ which classifies ChemOnt classes.^27^ Because NPClassifier classes are more appropriate for natural products than ChemOnt classes, BGC-MLM was compared to BGCat, although it should also be able to predict ChemOnt classes. BGCat is trained on BGCs from all sources, for a total 2,046 BGCs.^10^ Fine tuning for BGC-MLM was limited to BGCs from bacteria because it was only pretrained on bacterial BGCs. After eliminating BGCs for which SMILES strings were not available or for which NPClassifier did not make a prediction for, there were 1,224 training BGCs and 291 testing BGCs. BGCat predicts 294 classes, due to BGC-MLM’s smaller dataset several of these classes only had a few examples in the training set. Therefore, classification was limited to the seven NPClassifier pathways and 108 superclasses and classes that were present in more than three training examples and that are also predicted by BGCat. Due to the differences in training and test sets, and how many classes are predicted by each model, metrics are not directly comparable. By AUROC metrics, BGC-MLM models had similar performance (best scores of 0.95 micro averaged and 0.89 macro averaged, Table 1) compared to BGCat (best score 0.93 micro averaged and 0.81 macro averaged).^10^ BGC-MLM models had significantly higher recall (best score 0.90 micro averaged and 0.40 macro averaged) compared to BGCat (best score 0.44 micro averaged and 0.07 macro averaged) and significantly lower precision (best score 0.53 micro averaged and 0.26 macro averaged) compared to BGCat (best score 0.87 micro averaged and 0.82 macro averaged). This is likely because BGC-MLM was trained using class weighting, which penalizes the model more for false negatives from classes with low representation while BGCat does not. Class weighting leads to higher recall in exchange for lower precision.

**Table 1.** Performance on predicting NPClassifier classes. Average metrics across five models trained with different random seeds for each model type, standard deviation in parentheses. Best model in bold. AUROC calculation is limited to classes with positive labels for the test set. AP indicates average precision.

| Model | Micro Averaging |  |  |  | Macro Averaging |  |  |  |
| --- | --- | --- | --- | --- | --- | --- | --- | --- |
|  | AP | AUROC | Recall | Precision | AP | AUROC | Recall | Precision |
| Entire Test Set |  |  |  |  |  |  |  |  |
| Small frozen | 0.13 (0.01) | 0.93 (0.00) | **0.90 (0.01)** | 0.14 (0.01) | 0.09 (0.01) | 0.88 (0.00) | **0.40 (0.01)** | 0.10 (0.01) |
| Small finetune | 0.32 (0.02) | 0.94 (0.01) | 0.78 (0.01) | 0.40 (0.02) | 0.17 (0.01) | 0.87 (0.01) | 0.33 (0.01) | 0.20 (0.02) |
| Small scratch | 0.26 (0.02) | 0.94 (0.00) | 0.76 (0.01) | 0.34 (0.02) | 0.13 (0.01) | 0.88 (0.01) | 0.30 (0.01) | 0.16 (0.01) |
| Large frozen | 0.18 (0.01) | 0.94 (0.00) | 0.87 (0.01) | 0.21 (0.01) | 0.12 (0.01) | **0.89 (0.01)** | 0.38 (0.00) | 0.12 (0.01) |
| Large finetune | **0.39 (0.02)** | **0.95 (0.01)** | 0.73 (0.01) | **0.53 (0.01)** | **0.21 (0.01)** | 0.88 (0.01) | 0.30 (0.01) | **0.26 (0.01)** |
| Large scratch | 0.36 (0.02) | 0.93 (0.00) | 0.70 (0.01) | 0.50 (0.02) | 0.17 (0.01) | 0.87 (0.02) | 0.28 (0.01) | 0.22 (0.01) |
| Test Set with 20-40% Shared PFAMs with Finetuning Set |  |  |  |  |  |  |  |  |
| Small frozen | 0.10 (0.01) | 0.85 (0.01) | **0.80 (0.03)** | 0.12 (0.01) | 0.05 (0.00) | 0.84 (0.02) | **0.18 (0.01)** | **0.12 (0.01)** |
| Small finetune | 0.23 (0.04) | 0.87 (0.01) | 0.62 (0.04) | 0.37 (0.04) | **0.09 (0.01)** | 0.84 (0.01) | 0.13 (0.01) | 0.09 (0.01) |
| Small scratch | 0.16 (0.02) | 0.86 (0.01) | 0.56 (0.04) | 0.27 (0.03) | 0.07 (0.01) | 0.83 (0.01) | 0.11 (0.01) | 0.07 (0.01) |
| Large frozen | 0.15 (0.01) | 0.86 (0.01) | 0.76 (0.02) | 0.19 (0.01) | 0.07 (0.01) | 0.83 (0.02) | 0.16 (0.01) | 0.07 (0.01) |
| Large finetune | **0.27 (0.03)** | **0.88 (0.02)** | 0.53 (0.02) | **0.48 (0.04)** | 0.09 (0.00) | **0.85 (0.00)** | 0.11 (0.00) | 0.09 (0.01) |
| Large scratch | 0.24 (0.03) | 0.86 (0.01) | 0.54 (0.04) | 0.42 (0.03) | 0.08 (0.01) | 0.79 (0.01) | 0.10 (0.01) | 0.09 (0.01) |
| Test set with 20-40% Shared PFAMs with Pretraining Set |  |  |  |  |  |  |  |  |
| Small frozen | 0.08 (0.01) | 0.84 (0.01) | **0.77 (0.02)** | 0.10 (0.01) | 0.06 (0.00) | 0.83 (0.01) | **0.14 (0.01)** | 0.06 (0.01) |
| Small finetune | **0.18 (0.01)** | **0.88 (0.01)** | 0.63 (0.01) | 0.28 (0.02) | **0.07 (0.01)** | **0.85 (0.02)** | 0.11 (0.01) | **0.07 (0.01)** |
| Small scratch | 0.11 (0.00) | 0.86 (0.01) | 0.55 (0.03) | 0.19 (0.01) | 0.06 (0.00) | 0.83 (0.01) | 0.09 (0.01) | 0.06 (0.01) |
| Large frozen | 0.11 (0.01) | 0.84 (0.01) | 0.69 (0.04) | 0.15 (0.00) | 0.06 (0.00) | 0.81 (0.01) | 0.12 (0.01) | 0.06 (0.00) |
| Large finetune | 0.18 (0.02) | 0.87 (0.01) | 0.49 (0.02) | **0.34 (0.02)** | 0.06 (0.01) | 0.83 (0.02) | 0.07 (0.01) | 0.06 (0.01) |
| Large scratch | 0.15 (0.02) | 0.86 (0.01) | 0.47 (0.05) | 0.31 (0.02) | 0.06 (0.01) | 0.83 (0.03) | 0.07 (0.01) | 0.06 (0.01) |

Among BGC-MLM models, the large finetuned model performed the best (5/8 metrics), followed by the small frozen model (2/8) metrics (Table 1). The scratch models did not lead in any metrics, demonstrating the utility of pretraining. When models were evaluated only on the portion of the test set with 20-40% PFAMs shared with a finetuning or pretraining BGC, performance dropped considerably, especially for macro averaged metrics. However, the scratch models still did not perform best in any metrics, once again supporting the utility for pretraining for model generalizability.

### Prediction of Natural Product Bioactivity

We previously reported machine learning models for predicting bioactivity from protein annotations of BGCs, these models were based on relatively simplistic architectures (logistic regression, support vector machine classifiers, and extra random trees).^5^ Despite their simplicity, they have still proven to make correct predictions on BGCs not in their training set to guide the isolation of bioactive compounds.^28^ Due to its pretraining, BGC-MLM is expected to show improved accuracy and generalizability compared to these previous models. BGC-MLM was applied to the same training and hold out test datasets as our previously reported models.^5^ Our previous models were trained independently to predict each bioactivity. Because neural networks allow for multilabel classification, we trained one model that predicts all six activities we developed separate models for previously: antibacterial, anti-gram-positive bacteria, anti-gram-negative bacteria, anti-eukaryotic, antitumor/cytotoxic, and antifungal. To train this model, antibacterial compounds for which the target (gram-positive vs gram-negative) was not known were excluded. A separate model was trained for antibacterial, anti-eukaryotic, antitumor/cytotoxic, and antifungal activities with the antibacterial compounds of unknown bacterial target included.

Our previous models performed very poorly for portions of the holdout set that were distinct from the training set which included those that were not recognized by antiSMASH as BGCs as well as those with no or low (< 25%) knownclusterblast scores to training examples.^5^ This was especially true for the more difficult to predict bioactivity classes – anti-gram-negative and antifungal. Almost all BGC-MLM models outperformed our previous models at predicting anti-gram-negative and antifungal activities for the holdout sets BGCs unrecognized by antiSMASH, or with no or low (< 25%) knownclusterblast scores (Figure 3). BGC-MLM also usually outperformed previous models at prediction of other activities for BGCs in these segments of the holdout set (Figures 3, S5-10). The improved performance could be due to the larger number of parameters of BGC-MLM models compared to older models or the fact that BGC-MLM’s input allows it to learn from the order of domains, which older models could not do. Domain order, especially in the case of multimodular assembly line enzymes often define the product structure.^29^ Because domain order often determines structure and structure determines bioactivity, BGC-MLM having access to this information likely improves its performance.

**Figure 3.**
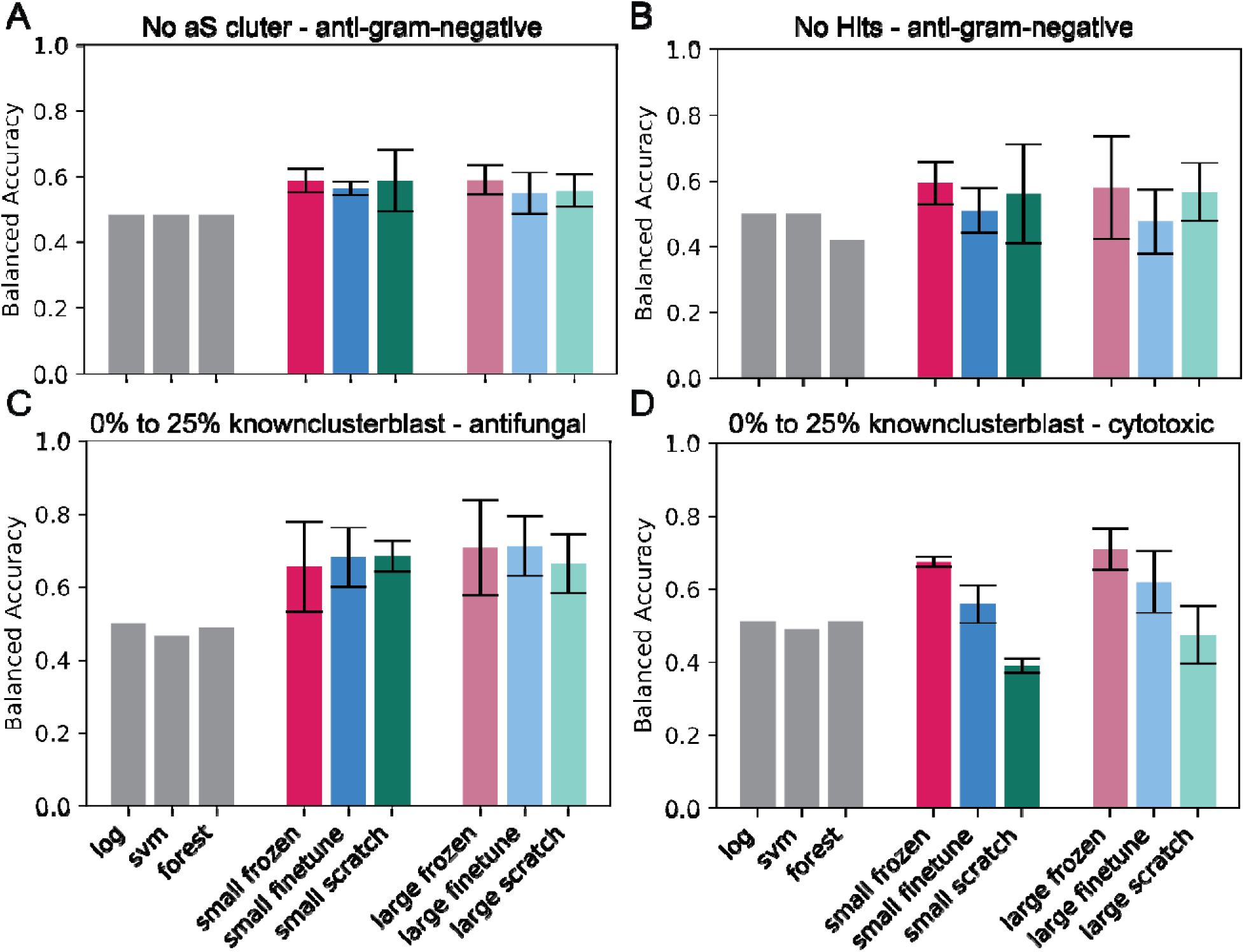
Prediction of bioactivities for splits of holdout set distinct from training data. Grey bars indicate predictions from our previous work (log = logistic regression, svm = support vector machine classifier, forest = extra random trees classifier), values are reproduced from this previous work.^5^ **A)** prediction of anti-gram-negative bacteria for BGCs not identified by antiSMASH. **B)** prediction of anti-gram-negative activity for BGCs with no hits by knownclusterblast **C)** prediction of antifungal activity for BGCs with knownclusterblast scores between 0% and 25%, models trained on dataset including BGCs without a known bacterial target **D)** prediction of cytotoxic/antitumor activity for BGCs with knownclusterblast scores between 0% and 25%, models trained on dataset including BGCs without a known bacterial target. Error bars for BGC-MLM models indicate standard deviation across five models trained with different random seeds. Only one model was trained for each type previously, so no error could be calculated for the previous models.

When applied to the whole holdout set, the different BGC-MLM models performed very similarly (Figure 4, S11). Based on the balanced accuracy metric, no scratch model was the best performing model for any bioactivity classification task. Therefore, like in the case of prediction of product compound class, pretraining improves performance, albeit only very slightly.

**Figure 4.**
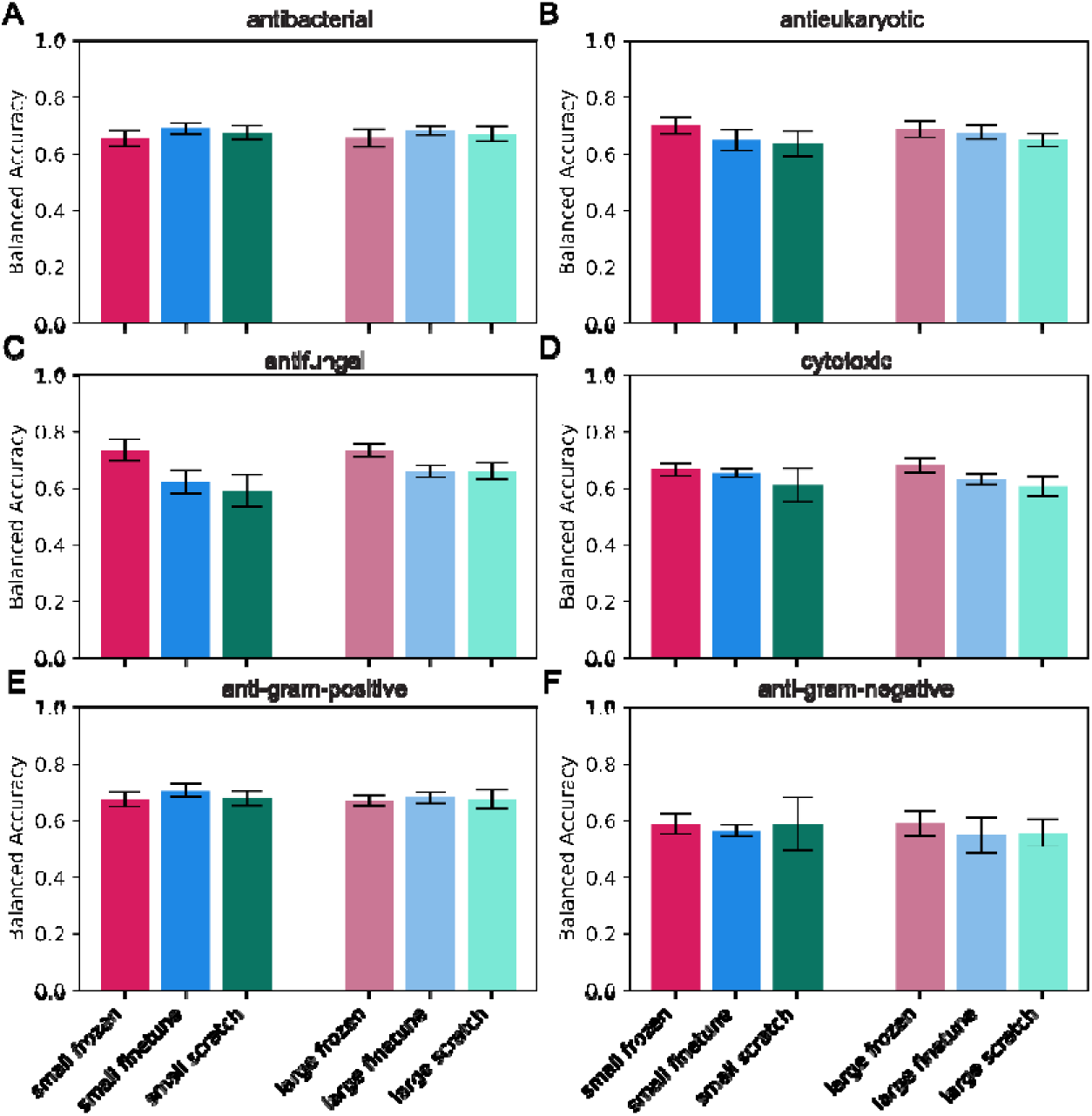
Prediction of bioactivity on entire holdout set. All models were trained on the training set that included classification for target for antibacterial activity. Error bars indicate standard deviation across five models trained wit different random seeds.

### Prediction of natural product molecular properties

The prediction of molecular properties of natural products from their BGCs is useful for genome mining and may be complementary to direct prediction of bioactivity because some molecular properties are known to be related to bioactivity. For example, amines are associated with the ability of molecules to accumulate in gram-negative bacteria and are therefore associated with activity against gram-negative bacteria.^30^ BGC-MLM was trained to predict general chemical properties (exact mass, total polar surface area, fraction of sp3 carbons, number of hydrogen bond acceptors, number of hydrogen bond donors, and calculated LogP) and functional groups or structures counts (aliphatic rings, aromatic rings, primary amines, carboxylic acids, amides, halogens, lactams, lactones, nitro groups, oxazoles, oximes, pyridines, thiazoles, and ureas). Prediction of these chemical descriptors can be done with a regression model. Because a BGC can produce more than one compound, the maximum and minimum value for each descriptor across all products for that BGC in the dataset were determined and treated as separate outputs. To build the regression model, the BGC-MLM embedding layers were connected to one fully connected layer, followed by an output layer. The model was then trained on the same stratified training set described above, eliminating any BGCs for which there were no SMILES string available through MIBiG or NPAtlas or that we had previously identified as having an error.^31^

BGC-MLM performed well at predicting some molecular properties including exact mass, total polar surface area, and number of H bond acceptors and donors (Pearson’s r > 0.6) (Figures 5, 6). It performed moderately well at predicting calculated LogP (cLogP) and poorly at predicting faction of sp3 carbons (Pearson’s r < 0.2). As seen with previous tasks, the scratch models were never the best performing and for the more difficult regression tasks performed notably worse, with Pearson’s r approaching zero for predicting fraction sp3 and cLogP. Performance was similar for predicting the maximum and minimum descriptor values across products of the same BGC (Figure S12).

**Figure 5.**
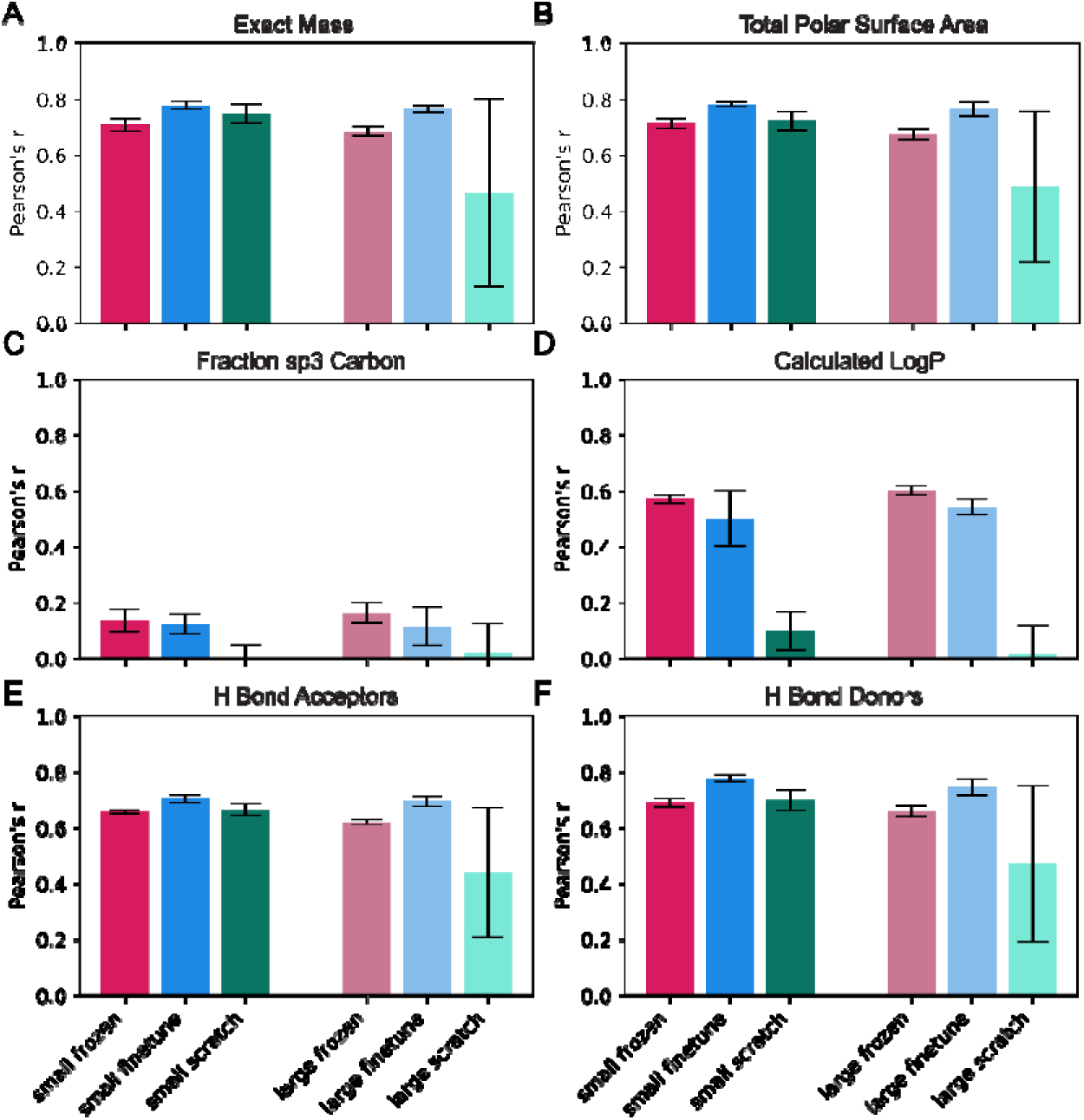
Prediction of maximum value of descriptors across BGC products. **A)** Exact mass **B)** total polar surface are **C)** fraction of sp3 carbons **D)** calculated LogP **E)** number of hydrogen bond acceptors **F)** number of hydrogen bond donors. Error bars indicate standard deviation across five models trained with different random seeds.

Performance at predicting counts of functional groups was more variable. The model’s performance for prediction of the number of amides was the best of all functional groups (Figures 6, 7), with Pearson’s r of 0.8 for the best performing model. Performance was very poor for some other functional groups such as oximes and ureas with Pearson’s r near zero. In general, performance was worse for less common functional groups and it is possible that these would fare better as classification tasks for presence/absence rather than regression tasks. The scratch models generally had worse performance than the pretrained models, again supporting that pretraining is an effective strategy. Performance was again similar for predicting the maximum and minimum number of functional groups over all products of the BGC (Figures 7, S13).

**Figure 6.**
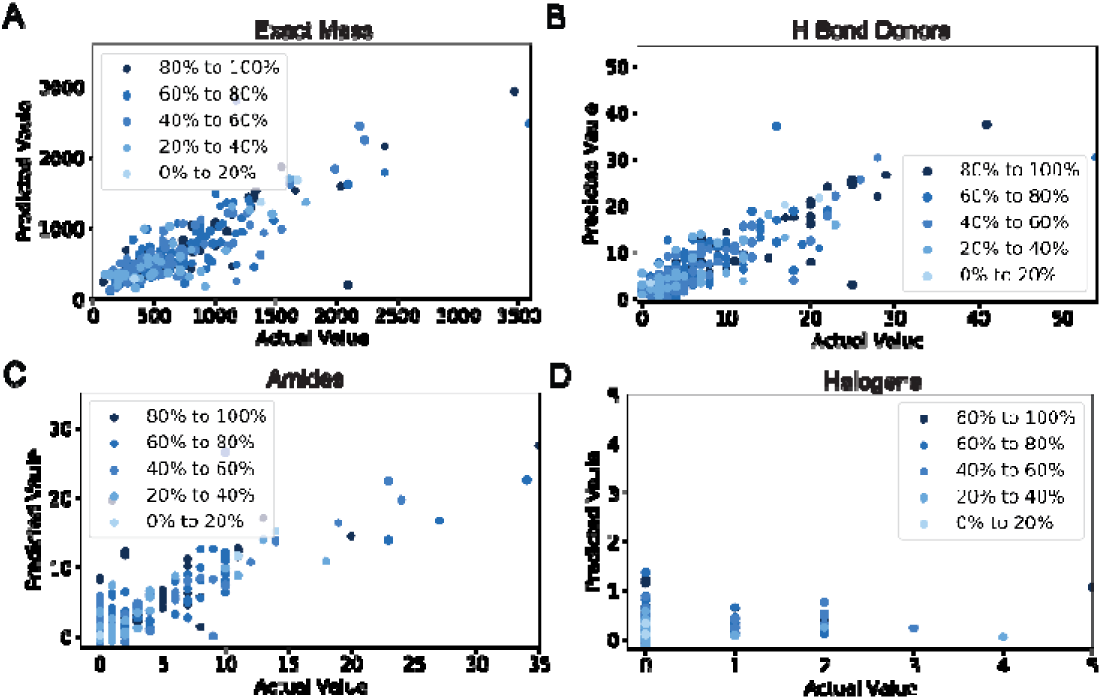
Actual properties vs predicted property for selected examples. **A)** exact mass **B)** number of hydrogen bon donors **C)** number of amides (good performance) **D)** number of halogens (poor performance). Dot color indicates maximum percentage shared PFAM domains with a finetuning BGC, with lighter dots indicating lower similarity.

**Figure 7.**
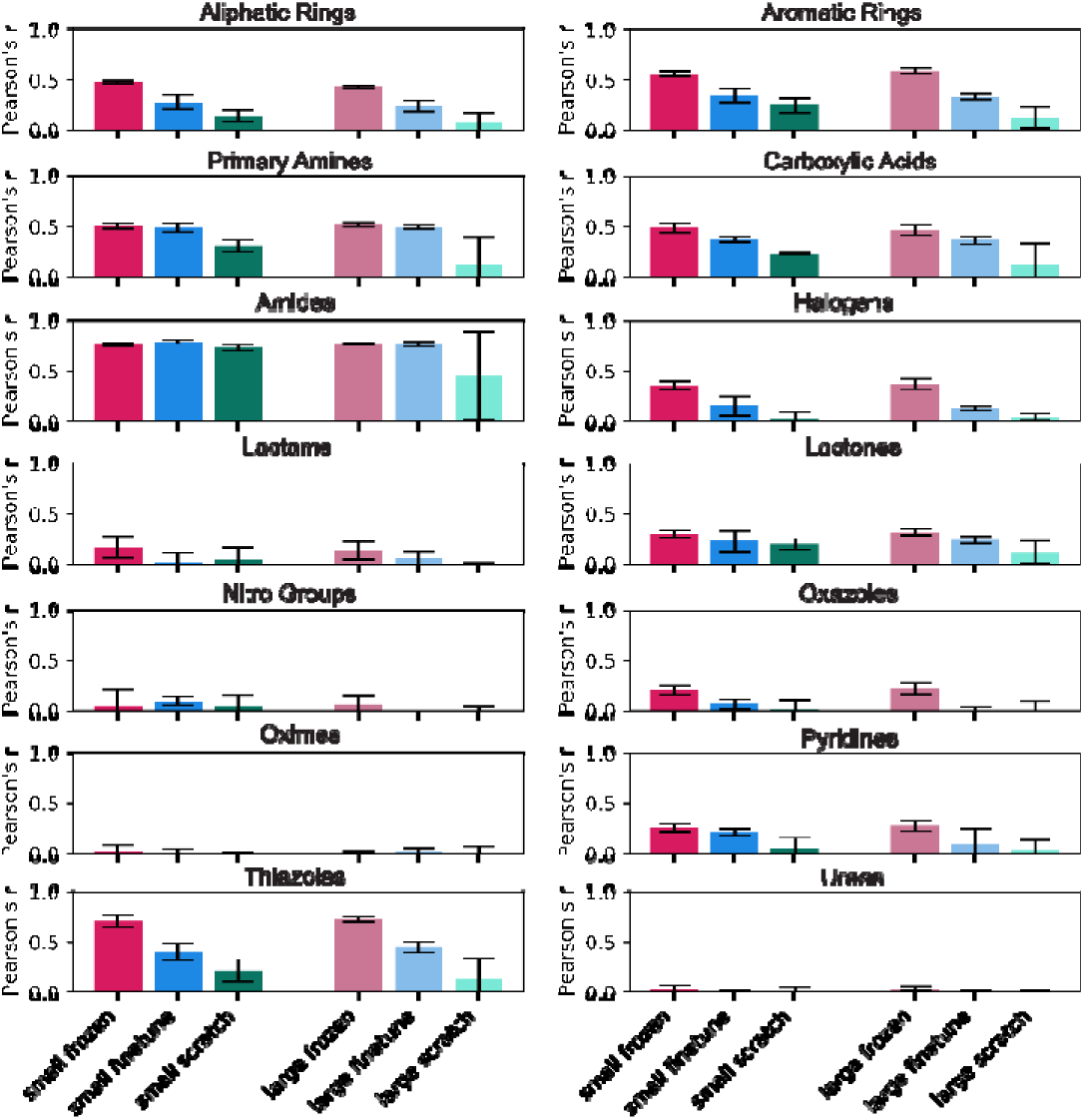
Prediction of maximum number of functional groups in products of a BGC. Bars indicate Pearson’ correlation coefficient. Error bars indicate standard deviation across five models trained with different random seeds.

### Prediction of Fingerprints and Linkage of BGCs and Products

Prediction of molecular fingerprints, bit vectors indicating the presence or absence of substructures where either defined substructure indices or hash functions of the substructure determine the associated bit in the vector,^32,33^ can be treated as a multilabel classification problem. Prediction of fingerprints allows for comparison of predicted fingerprints to actual fingerprints of natural products. The similarity of predicted fingerprints to actual fingerprints allows for linking BGCs to compounds and vice versa by ranking BGC-molecule pairs by the similarity of the predicted and actual fingerprints. There are multiple different types of molecular fingerprints and many of these fingerprints can also have different sizes. Larger fingerprints reduce the chance of collisions which is when two substructures are represented by the same bit. Fingerprints can also be compared using cosine similarity or Tanimoto similarity.^34^ To determine which fingerprint type, size, and comparison method were optimal for BGC-product linkage, a small frozen model was trained to predict each fingerprint supported by RDKit at different sizes, except for MACCS fingerprints, which have a fixed size. These models were also trained with and without class weights. The models were then used to predict fingerprints for the stratified test set described above. To test the ability of the predicted fingerprints to link BGC and product, the average rank of the correct BGC among all BGCs in training and test sets and the average rank of the first correct molecule among all products of BGCs in the training and test sets were determined. Models without class weighting performed better than those with class weighting. Morgan fingerprints with a size of 2048 performed the best at linking BGC to molecule, while MACCS performed the best at the reverse problem (Tables S5, S6).

Five models were then trained for each model type and for Morgan fingerprints of size 2048 and MACCs fingerprints. Based on most micro and macro averaged metrics, the large finetuned model performed the best for both fingerprints, with metrics for MACCS being much higher, likely because there are fewer rare bits because the MACCS fingerprint is shorter than the 2048 size used for the Morgan fingerprints (Table S7).

The models for MACCS fingerprint prediction were then investigated for their ability to link molecule to BGC for only the genome of the producing organism. This is a common problem in natural product discovery, where the producing strain and compound is known and researchers want to determine which BGC is likely responsible for production of the natural product. This task is typically accomplished by manual inspection of BGCs. Models that link products to their BGC can make this process more efficient because if they are accurate, researchers will only need to inspect the top BGCs chosen by the model to identify the BGC responsible for producing the compound. Because cosine similarity worked better than Tanimoto similarity across all BGCs and molecules (Table S6), cosine similarity was used for this task. Ten test set molecules produced by eight unique BGCs from different biosynthetic classes were chosen as case studies. If the models have no predictive utility, it is expected that the correct BGC will be ranked at around half the total number of BGCs on average. Only one of the chosen molecules, catenulipeptin, showed performance near what was expected for a model with no predictive utility and all other compounds showed better performance (Table 2). The poor performance on catenulipeptin can likely be explained by the fact that it is a lanthipeptide and there are multiple lanthipeptide BGCs encoded in the same genome. Lanthipeptide structure is primarily determined by the sequence of the core region of the precursor peptide which BGC-MLM does not have access to, making it difficult for BGC-MLM to correctly predict the fingerprint of lanthipeptides and likely other RiPPs. For five of the compounds, from three unique BGCs, at least one BGC-MLM model was able to correctly identify the producing BGC. Once again, the models trained without pretraining either matched or performed worse than other models, demonstrating the advantage of pretraining.

**Table 2.**
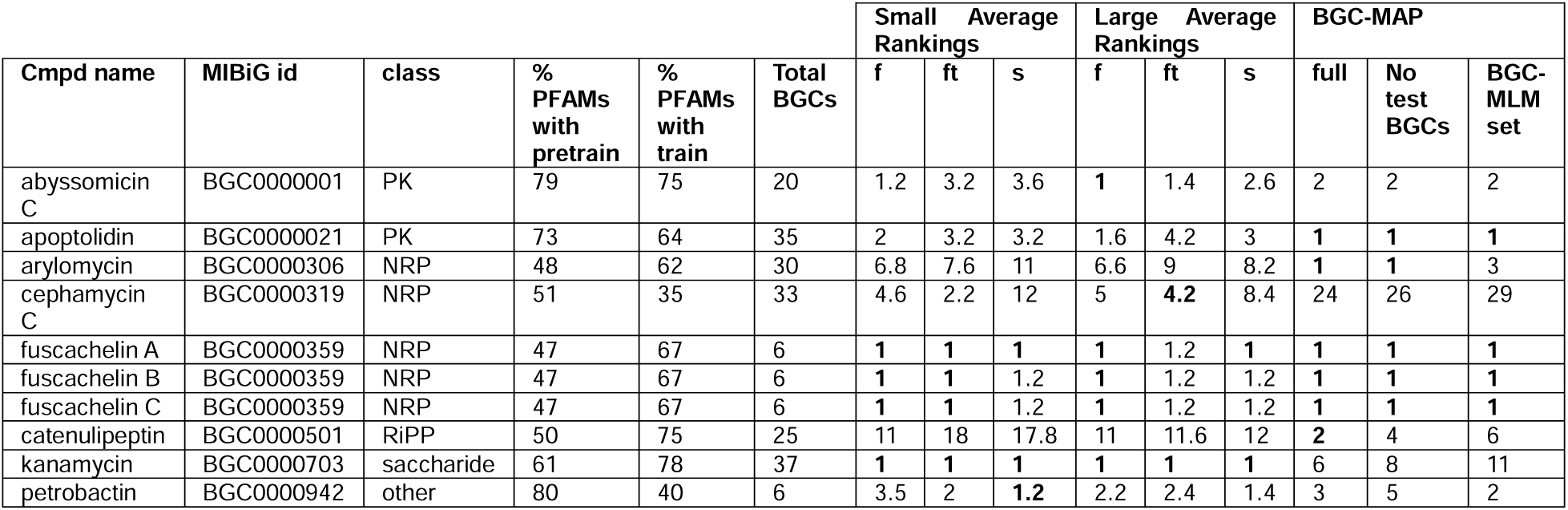
Case studies in linking molecule to BGC with BGC-MLM and BGC-MAP. Bold indicates best performance for that compound.

BGC-MLM was compared to another model for ranking BGCs by likelihood of producing a product, BGC-MAP.^22^ BGC-MAP performed better than the best BGC-MLM model for three of the ten compounds and tied BGC-MLM’s performance for the three fuscachelin congeners (Table 2). When BGC-MAP was retrained with this test set examples removed from its training set or with the same set that was used for fine-tuning BGC-MLM, performance on prediction of the arylomycin and catenulipeptin prediction dropped but remained higher than all BGC-MLM models. Interestingly, for two of the molecules BGC-MLM performed relatively well on, cephamycin C and kanamycin, BGC-MAP performed much worse even when these compounds. Overall, BGC-MLM’s performance is comparable to BGC-MAP with better performance in some cases and worse in others.

### Comparison and Clustering of BGCs with Fingerprint Prediction

Previously, we developed a set of benchmarks to assess methods for comparing and clustering BGCs on how well these comparisons correspond to structural similarity of their products.^31^ In this benchmark, BiG-SCAPE was generally the best performing method of those that perform both similarity calculation and clustering. After our benchmark study BiG-SCAPE 2 was released,^35^ so to determine an ideal comparison BiG-SCAPE 2 was benchmarked. BiG-SCAPE 2 had a similar correlation to BiG-SCAPE between distance or score, respectively, and Tanimoto similarity (−0.25 vs 0.26), given the limited improvement of BiG-SCAPE 2 by this metric BGC-MLM was compared to the original best results from our previous benchmarking paper. To compare similarity between two BGCs, BGC-MLM can be used to predict the fingerprint and the Tanimoto similarity between predicted fingerprints can be calculated. The best correlation between predicted similarity and Tanimoto similarity of the product using this method was 0.50 (Table 3). The benchmark set overlaps with our training set, so correlation with training data removed from the benchmark dataset was also calculated. This decreased performance, with the best model having a correlation of 0.33, however this correlation still exceeds the best performance on methods when they are forced to calculate similarities across all pairs of BGCs (best performance is clust-o-matic with a correlation of 0.28) and is not dramatically lower than the overall best performing method, knownclusterblast with a correlation 0.47.^31^ Once again, the scratch models were not the best model, demonstrating the advantage of pretraining.

**Table 3.**
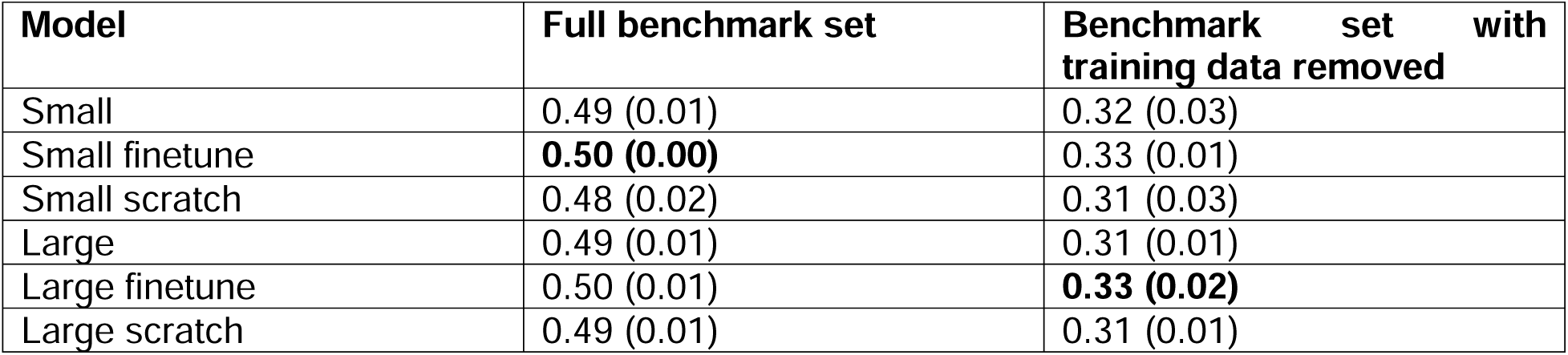
Spearman’s correlation of BGC-MLM predicted fingerprint similarity and product Tanimoto similarity. Average across five models, standard deviation in parentheses.

| Model | Full benchmark set | Benchmark set with training data removed |
| --- | --- | --- |
| Small | 0.49 (0.01) | 0.32 (0.03) |
| Small finetune | **0.50 (0.00)** | 0.33 (0.01) |
| Small scratch | 0.48 (0.02) | 0.31 (0.03) |
| Large | 0.49 (0.01) | 0.31 (0.01) |
| Large finetune | 0.50 (0.01) | **0.33 (0.02)** |
| Large scratch | 0.49 (0.01) | 0.31 (0.01) |

**Table 4.** BGC-MLM predicted fingerprints clustering adjusted Rand scores. Average across five models, standard deviation in parentheses.

| Model | Full benchmark set | Benchmark set with training data removed |
| --- | --- | --- |
| Small | 0.20 (0.02) | 0.09 (0.03) |
| Small finetune | 0.25 (0.01) | 0.16 (0.04) |
| Small scratch | 0.20 (0.02) | 0.08 (0.04) |
| Large | 0.21 (0.03) | 0.10 (0.05) |
| Large finetune | 0.24 (0.01) | 0.13 (0.06) |
| Large scratch | 0.18 (0.02) | 0.06 (0.02) |
| Best of methods compared in previous study<sup>31</sup> | 0.66 | NA |

The predicted fingerprints can also be clustered to cluster BGCs into groups that produce compounds with similar structures. The Tanimoto similarity of the predicted fingerprints was calculated, and the OPTICS clustering method was used to cluster the BGCs. Using this methodology, none of the BGC-MLM models were able to achieve the performance of existing methods. For example, the adjusted Rand metric comparing BGC clusters and clusters of molecular structure calculated with the Butina clustering method was between 0.18 and 0.25 across all BGC-MLM models for the full benchmark set compared to a best performance of 0.66 across existing methods. Given that BGC-MLM is more accurate at calculating BGC similarity in a way that reflects structural similarity, it is likely the clustering method that is responsible for the lower performance and additional optimization of clustering is required to make clustering of BGC-MLM predicted fingerprints competitive with existing methods. The models trained without pretraining had the worst performance, once again highlighting the benefits of pretraining.

## Conclusion

BGC-MLM is a pretrained foundation model that can be used to predict multiple properties of the products of BGCs. Specifically, it can predict both broad and fine-grained structural class, bioactivity, molecular properties, counts of functional groups, and molecular fingerprint. Prediction of molecular fingerprints can be used to link BGCs to molecules and vice versa and to compare BGC similarity and cluster BGCs by product similarity. Across all applications, models that were pretrained outperformed those that were not, showing that pretraining with a masked language task is a useful approach for building foundation models for BGCs. BGC-MLM also demonstrates comparable or better performance than specialized methods for all tasks except clustering BGCs by predicted product similarity, demonstrating its utility as a foundation model. BGC-MLM will enable the prioritization of BGCs predicted to produce compounds with desirable bioactivities, molecular properties, and substructures/functional groups. It can also be adapted for many other BGC-related tasks. One possible future application would be to directly predict natural product structure with a sequence-to-sequence model, however given the variable performance of BGC-MLM on prediction of structural properties, additional pretraining and fine-tuning data or further architectural improvements are likely needed before accurate natural product structure prediction can be realized. Overall, BGC-MLM represents a promising foundation model for BGCs that can be used to guide genome mining of products with desirable bioactivities or molecular properties.

## Methods

### Pretraining Dataset construction

The pretraining dataset was constructed from a dataset of BGCs predicted using antiSMASH 5 with PFAM detection turned on.^23^ Part of this dataset was reported in our previous publication^24^ and was expanded in this work. BGCs identified as being on a contig edge by antiSMASH were excluded, leaving a total of 1,061,133 BGCs. The PFAM domains with scores greater than or equal to 20 were extracted from the antiSMASH output genbank files using a custom python script tokenize_bgcs_pfam_only.py.

### Pretraining training and test set split

To avoid inflated test accuracies from data leakage, a maximum number of shared PFAM annotations at 70% for pairs of data in the training and test set was set – meaning that for a BGC in the test set, all BGCs in the training set will share less than 70% of the PFAM annotations with that BGC. The percentage of PFAMs shared was calculated by the following formula

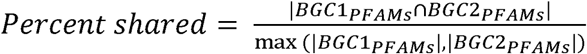

where BGC1PFAMs and BGC2PFAMS are the set of PFAMS in BGC1 and BGC2, respectively. To accomplish this split, data was first split randomly with 90% assigned to the training and the remainder in the test data. All BGCs originally assigned to test set that had greater than 70% similarity with a BGC in the training set was moved to the training set. This process was repeated, comparing the remaining test data to the recently moved BGCs and until there were no test BGCs that needed to be moved to the training dataset. After this process was completed, there were 994,418 training and 5,582 test BGCs. Because many of the test BGCs had been moved, those that remained were likely unusual, very rare, or even false positive BGCs. To ensure the test set had some more standard BGCs, an additional 20 random BGCs from the new training were moved to the test set, and then all BGCs with over 70% of shared PFAMs to those BGCs were also moved to the test set. This process was repeated for the recently moved BGCs until there were no remaining BGCs to move. At the end of this process there were 897,008 training and 164,125 test BGCs. The full pretraining and corresponding test set is available at https://huggingface.co/datasets/allie-walker/BGC_MLM_data.

### Pretraining

Pretraining was done using a masked language model (MLM), similar to that reported by Devlin et al.^13^ but without the sentence pair identification component. Specifically, the implementation by CheeKean^36^ was used as a starting point to write the MLM code. BGCs of length greater than 100 were excluded and the remaining BGCs were padded with a PAD token to make all BGCs the same length. PFAMs that occurred fewer than 20 times in the training set were replaced with an unknown “<UNK>” token. 15% of tokens are selected for masking and of those 80% are masked, 10% are changed to a different token, and 10% remain unchanged. The model is made up of an embedding layer followed by encoding blocks consisting of a layer normalizer, multiheaded attention, a feed forward layer, and dropout. The number of encoder blocks, the hidden representation size, the number of attention heads, and dropout rate can be varied. The script used to pretrain the models is BGC_MLM_train.py.

### Analysis of pretrained predictions

Pretrained models were assessed using accuracy and top-K accuracy with K of five or ten. The script used to calculate these metrics is topKPrediction.py.

### Fine-tuning for multi-label classification tasks

Natural product class, molecular fingerprint, and bioactivity prediction tasks were all treated as multi-label classification tasks because in all cases it is possible for a single BGC to have multiple labels. The classifier architecture consisted of BGC-MLM, where the final layer before the output layer was combined by mean pooling, followed by one feed forward layer, and then one output layer for classification. BCEWithLogitsLoss was used as the loss function. In all cases three versions of each model size were trained, one with all weights from the pretrained model frozen one with all pretrained weights unfrozen, and a model trained from scratch with no pretrained weights. The classificationTask.py script is used to train the frozen and fine-tuned models and the classificationTaskFromScratch.py script is used to train the scratch models. Performance metrics were calculated using scikit-learn and the test set was further divided by percent PFAMs shared with the pretraining or fine-tuning datasets for all tasks except bioactivity prediction. For bioactivity prediction, the test set was further divided by knownclusterblast score as described previously.^5^ Training and test data for all tasks is available on our HuggingFace repository and additional details on dataset construction and training for each task are described below.

### MIBiG class

Genbank and JSON files associated with bacterial BGCs were downloaded from MIBiG v4.^25^ Natural product class was extracted from the JSON file using the custom script getClassificationsFromJSON.py. The training and test set were split based on a stratified split on product class with 20% assigned to the test set. The multi-label classifier model described above was trained to predict product class. Models were trained for 75 epochs with 90% of the data used as training data and the remaining 10% as validation data. Training was run with class weighting.

### NPClassifier class

NPClassifier classifications were obtained using a custom script run_NPClassifier_API.py which obtains classifications using the NPClassifier API. NPClassifier did not predict a pathway for a molecule it was removed from the dataset. As described for BGCat, if a BGC had multiple products, their classifications were combined for the multilabel classification task.^10^ The same training and testing split as for the MIBiG classification tasks were used. The models were trained for 100 epochs with 99% of data used as training and 1% used as validation. Training was run with class weighting.

### Natural product bioactivity

Previously reported training and holdout test datasets^5^ were used to train the natural product bioactivity classifiers. The models were trained for 150 epochs with 99% of data used as training and 1% used as validation. Training was run with class weighting.

### Natural product fingerprint

To construct a fine-tuning training set for fingerprint prediction, it was necessary to first construct a dataset linking BGC to product structure. Bacterial BGCs from the MIBiG database^25^ and a modified version of the NPAtlas database^37^ reported in our BGC similarity benchmarking study,^31^ which was corrected to remove BGC-product linkages that were incorrect and add missing BGC-product linkages was used as a starting point. Then, linkages from MIBiG to NPAtlas compounds that were not in NPAtlas were added by extracting these linkages from MIBiG JSON files. BGCs we previously identified as having errors were excluded.^31^ This dataset will be referred to as the MIBiG-NPAtlas dataset.

Fingerprints for each compound were generated by RDKit, specifically the atom pair, functional Morgan, Morgan, RdKit, and topological torsion fingerprints. Each of these fingerprints were tested with the following sizes: 2048, 4096, and 8192. MACCS fingerprints which have a fixed size were also tested. For BGCs that are associated with multiple products, fingerprints were combined by taking the union of the two fingerprints, such that if at least one fingerprint for a product of a BGC had a specific bit, it would be present in the combined fingerprint. The multi-label classifier described above was then trained to predict fingerprints. The models were trained for 200 epochs with 90% of the data used for training and 10% for validation. Training was run both with and without class weighting.

To assess the fingerprint prediction models, the models were used to rank associations between BGCs and molecules and vice versa. To test the ability of the model to link BGC to molecule, each fingerprint predicted for the class-stratified test set with an associated product, was compared to the fingerprints of all molecules in the MIBiG-NPAtlas dataset using cosine and Tanimoto similarity scores. These compounds were then ranked by each similarity score and the rank of the highest-ranking true product of the BGC was determined. To test the ability of the model to link a molecule to its BGC, all molecules associated with the class-stratified test set were compared to all predicted fingerprints from the MIBiG-NPAtlas dataset using cosine and Tanimoto similarity and BGCs were ranked by similarity to the molecular fingerprint. The rank of the correct BGC was then determined. The fpRankMetrics.py script was used to perform this ranking analysis.

A similar analysis was performed for linking molecules to BGCs but limited to BGCs from the genome of a known producer of the molecule. antiSMASH 5 was first used to predict the BGCs from the genome, the BGCs were tokenized as described above and the fpRankGenome.py script was used to determine the ranking of each BGC based on the predicted fingerprint.

For comparison of predicted fingerprint similarity to BGC similarity, fingerprints were predicted for each BGC in our previously published benchmark dataset,^31^ MACCs fingerprints were predicted, and Tanimoto similarities were calculated for all pairs and our previous benchmarking scripts were used to calculate Spearman’s correlation between predicted fingerprint’s Tanimoto similarity and Tanimoto similarity of the products. The pairwise Tanimoto similarity matrix of the predicted fingerprints was also used to cluster BGCs using the OPTICS clustering algorithm and our previously published benchmarking scripts were used to determine cluster quality compared to the products clustered by the Butina clustering algorithm.^38^

### Fine-tuning to predict molecular properties and functional groups

RDKit was used to calculate molecular properties and the number of different functional groups, also referred to as descriptors, present in all molecules in the MIBiG-NPAtlas dataset. Specifically, the following descriptors were calculated: exact mass, total polar surface area, fraction of sp3 carbons, number of hydrogen bond acceptors, number of hydrogen bond donors, calculated LogP, number of aliphatic rings, number of aromatic rings, number of primary amines, number of carboxylic acids, number of amides, number of halogens, number of lactams, number of lactones, number of nitro groups, number of oxazoles, number of oximes, number of pyridines, number of thiazoles, number of ureas. The minimum and maximum values for each descriptor for each BGC across all products associated with that BGC were determined. A regression model was then trained on stratified training dataset for which structures were available. The regression model followed the same architecture as the previously described multilabel classifier except with MSELoss used as the loss function. As with the classification task, models were trained with frozen pretrained weights, unfrozen pretrained weights, and from scratch. The model can be trained with regressionTask.py for frozen and fine-tuned models and regressionTaskFromScatch.py for the scratch models. The accuracy of the model was assessed using Mean Absolute Error (MAE) and Pearson’s r on the stratified test set and on different subsets of the stratified test set split by similarity to the pretraining or fine-tuning set.

### Data and code availability

All code is available at https://github.com/aswalker-lab/BGC-MLM, all data at https://huggingface.co/datasets/allie-walker/BGC_MLM_data, and all model weights at https://huggingface.co/allie-walker/BGC_MLM.

## Acknowledgments

Research reported in this publication was supported by the National Institute of General Medical Sciences under award number R35GM146987. The content is solely the responsibility of the author and does not necessarily represent the official views of the National Institutes of Health. Computational resources were provided by Vanderbilt’s ACCRE and Center for Structural Biology.

