## Supplemental information for "A foundation model enables prediction of natural product molecular properties, bioactivity, and structural similarity from biosynthetic gene cluster sequence"

**Table S1. Hyperparameter optimization results**

| **Hidden dimension** | **Number of layers** | **Number of heads** | **Dropout** | **Train top 1 accuracy** | **Train top 10 accuracy** | **Test top 1 accuracy** | **Test top 10 accuracy** |
| --- | --- | --- | --- | --- | --- | --- | --- |
| 768 | 2 | 12 | 0.1 | 0.95 | 0.98 | 0.95 | 0.99 |
| 1536 | 2 | 12 | 0.1 | 0.96 | 0.99 | 0.97 | 0.99 |
| 768 | 3 | 12 | 0.1 | 0.95 | 0.98 | 0.96 | 0.99 |
| 896 | 2 | 14 | 0.1 | 0.95 | 0.98 | 0.95 | 0.99 |
| 1792 | 2 | 14 | 0.1 | 0.96 | 0.99 | 0.96 | 0.99 |
| 768 | 2 | 12 | 0.5 | 0.93 | 0.97 | 0.95 | 0.99 |
| 768 | 2 | 12 | 0.05 | 0.95 | 0.98 | 0.96 | 0.99 |
| 3072 | 2 | 12 | 0.1 | 0.96 | 0.99 | 0.96 | 0.99 |
| 640 | 2 | 10 | 0.1 | 0.95 | 0.98 | 0.96 | 0.99 |

**Table S2. Independently trained small (hidden dimension 768) and large (hidden dimension 1792) models**

| **Model type** | **Train top 1 accuracy** | **Train top 10 accuracy** | **Test top 1 accuracy** | **Test top 10 accuracy** |
| --- | --- | --- | --- | --- |
| Small | 0.95 | 0.98 | 0.96 | 0.99 |
| Small | 0.95 | 0.98 | 0.96 | 0.99 |
| Small | 0.95 | 0.98 | 0.96 | 0.99 |
| Small | 0.95 | 0.98 | 0.96 | 0.99 |
| Large | 0.96 | 0.99 | 0.97 | 0.99 |
| Large | 0.96 | 0.99 | 0.97 | 0.99 |
| Large | 0.96 | 0.99 | 0.97 | 0.99 |
| Large | 0.96 | 0.99 | 0.97 | 0.99 |

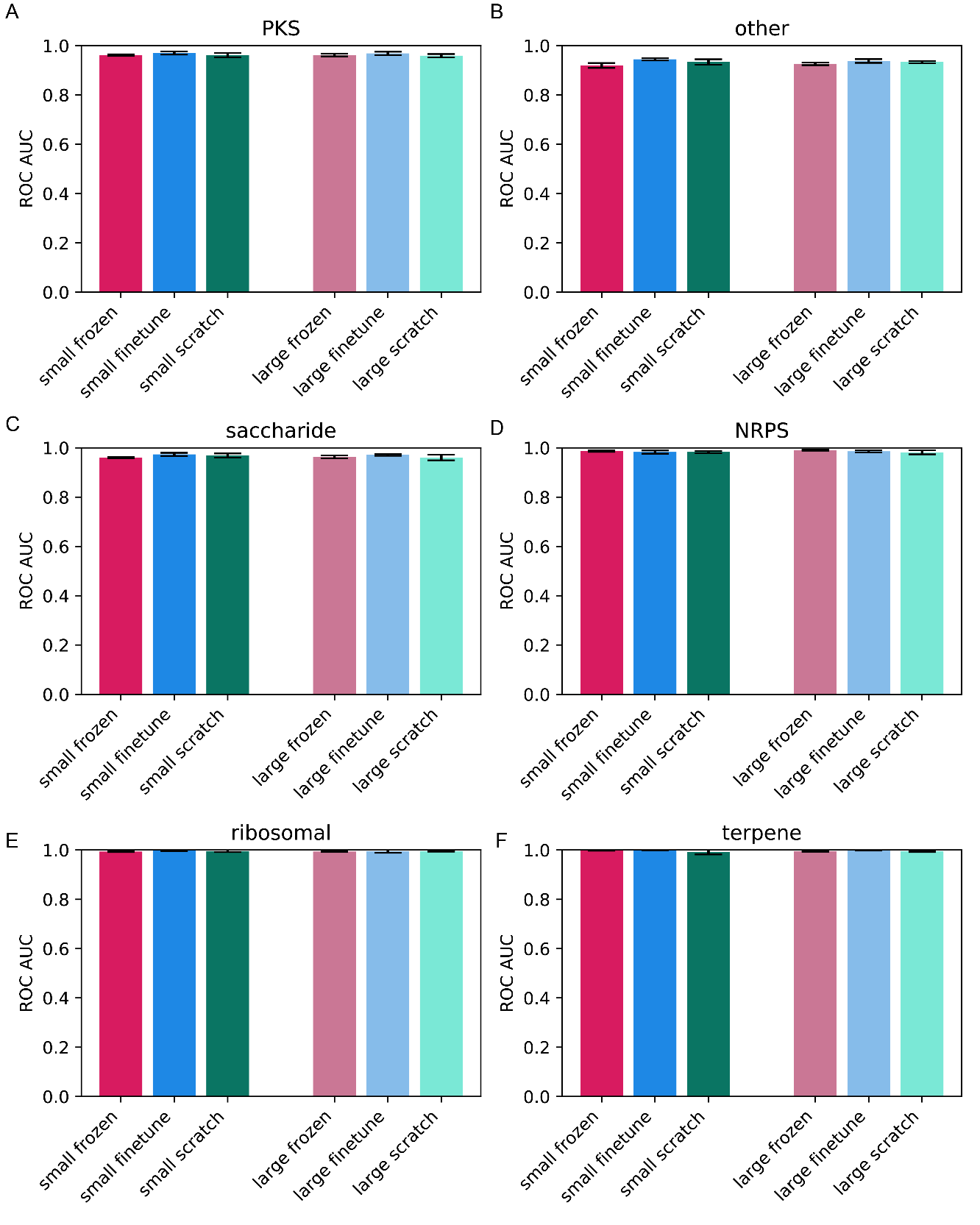

**Figure S1. AUROC for MIBiG classifications, full test set.** Error bars indicate standard deviation across five models.

**Table S3. Average performance on MIBiG classifications, full test set.** Best performing model for class in bold (based on unrounded values in case of tie). Values that round to 1.00 are indicated as “>0.99” to avoid the appearance of perfect performance.

| **Model** | **AUC ROC** | **AUC PRC** |
| --- | --- | --- |
| **PKS** | | |
| Small frozen | 0.96 | 0.93 |
| Small finetune | **0.97** | **0.94** |
| Small scratch | 0.96 | 0.93 |
| Large frozen | 0.96 | 0.93 |
| Large finetune | 0.97 | 0.94 |
| Large scratch | 0.96 | 0.92 |
| **other** | | |
| Small frozen | 0.92 | 0.74 |
| Small finetune | **0.94** | 0.81 |
| Small scratch | 0.93 | **0.81** |
| Large frozen | 0.94 | 0.77 |
| Large finetune | 0.93 | 0.80 |
| Large scratch | 0.93 | 0.80 |
| **NRPS** | | |
| Small frozen | 0.99 | 0.98 |
| Small finetune | 0.98 | 0.98 |
| Small scratch | 0.98 | 0.98 |
| Large frozen | **0.99** | **0.98** |
| Large finetune | 0.99 | 0.98 |
| Large scratch | 0.98 | 0.98 |
| **ribosomal** | | |
| Small frozen | 0.99 | 0.98 |
| Small finetune | **>0.99** | **0.99** |
| Small scratch | >0.99 | 0.99 |
| Large frozen | 0.99 | 0.98 |
| Large finetune | 0.99 | 0.99 |
| Large scratch | >0.99 | 0.99 |
| **saccharide** | | |
| Small frozen | 0.96 | 0.78 |
| Small finetune | **0.97** | 0.86 |
| Small scratch | 0.97 | 0.85 |
| Large frozen | 0.96 | 0.82 |
| Large finetune | 0.97 | **0.87** |
| Large scratch | 0.96 | 0.84 |
| **terpene** | | |
| Small frozen | >0.99 | 0.95 |
| Small finetune | **>0.99** | **0.97** |
| Small scratch | 0.99 | 0.90 |
| Large frozen | >0.99 | 0.93 |
| Large finetune | >0.99 | 0.96 |
| Large scratch | 0.99 | 0.90 |

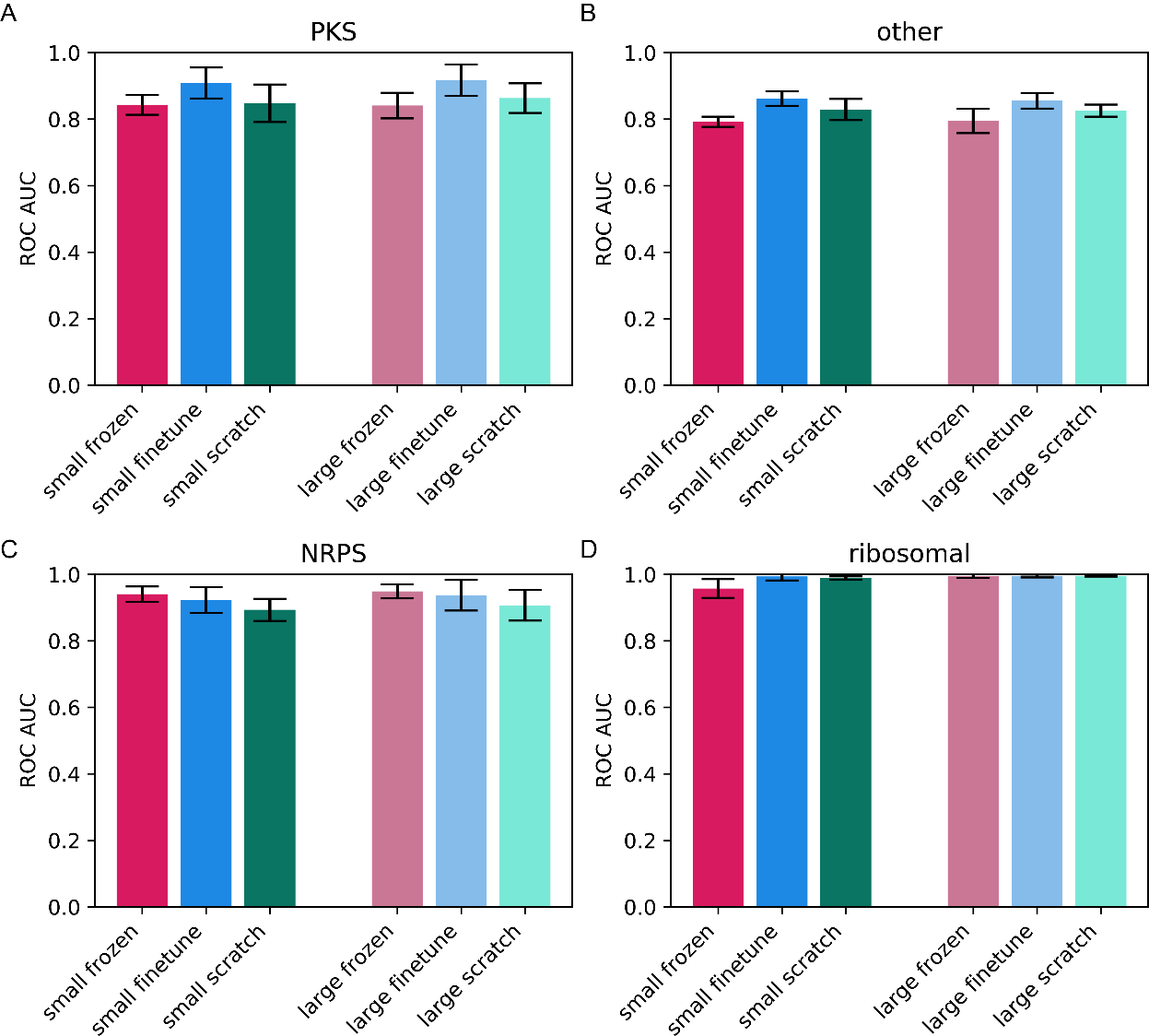

**Figure S2. AUROC for MIBiG classifications, test set with 20-40% similarity to fine-tuning set.** Classes that are not shown did not have enough BGCs in the 20-40% similarity range to calculate metrics. Error bars indicate standard deviation between models trained with different random seeds.

**Table S4. Average performance on MIBiG classifications, 20-40% similarity to fine-tune set.** Best performing model for class in bold (based on unrounded values in case of tie). Values that round to 1.00 are indicated as “>0.99” to avoid the appearance of perfect performance.

| **Model** | **AUC ROC** | **AUC PRC** |
| --- | --- | --- |
| **PKS** | | |
| Small frozen | 0.84 | 0.58 |
| Small finetune | 0.91 | **0.74** |
| Small scratch | 0.85 | 0.58 |
| Large frozen | 0.84 | 0.64 |
| Large finetune | **0.92** | 0.73 |
| Large scratch | 0.86 | 0.62 |
| **other** | | |
| Small frozen | 0.79 | 0.77 |
| Small finetune | **0.86** | 0.83 |
| Small scratch | 0.83 | 0.83 |
| Large frozen | 0.79 | 0.78 |
| Large finetune | 0.86 | **0.85** |
| Large scratch | 0.83 | 0.81 |
| **NRPS** | | |
| Small frozen | 0.94 | 0.88 |
| Small finetune | 0.92 | 0.86 |
| Small scratch | 0.89 | 0.85 |
| Large frozen | **0.95** | **0.89** |
| Large finetune | 0.94 | 0.89 |
| Large scratch | 0.91 | 0.86 |
| **ribosomal** | | |
| Small frozen | 0.96 | 0.89 |
| Small finetune | 0.99 | 0.98 |
| Small scratch | 0.99 | 0.95 |
| Large frozen | >0.99 | 0.98 |
| Large finetune | >0.99 | 0.98 |
| Large scratch | **>0.99** | **0.98** |

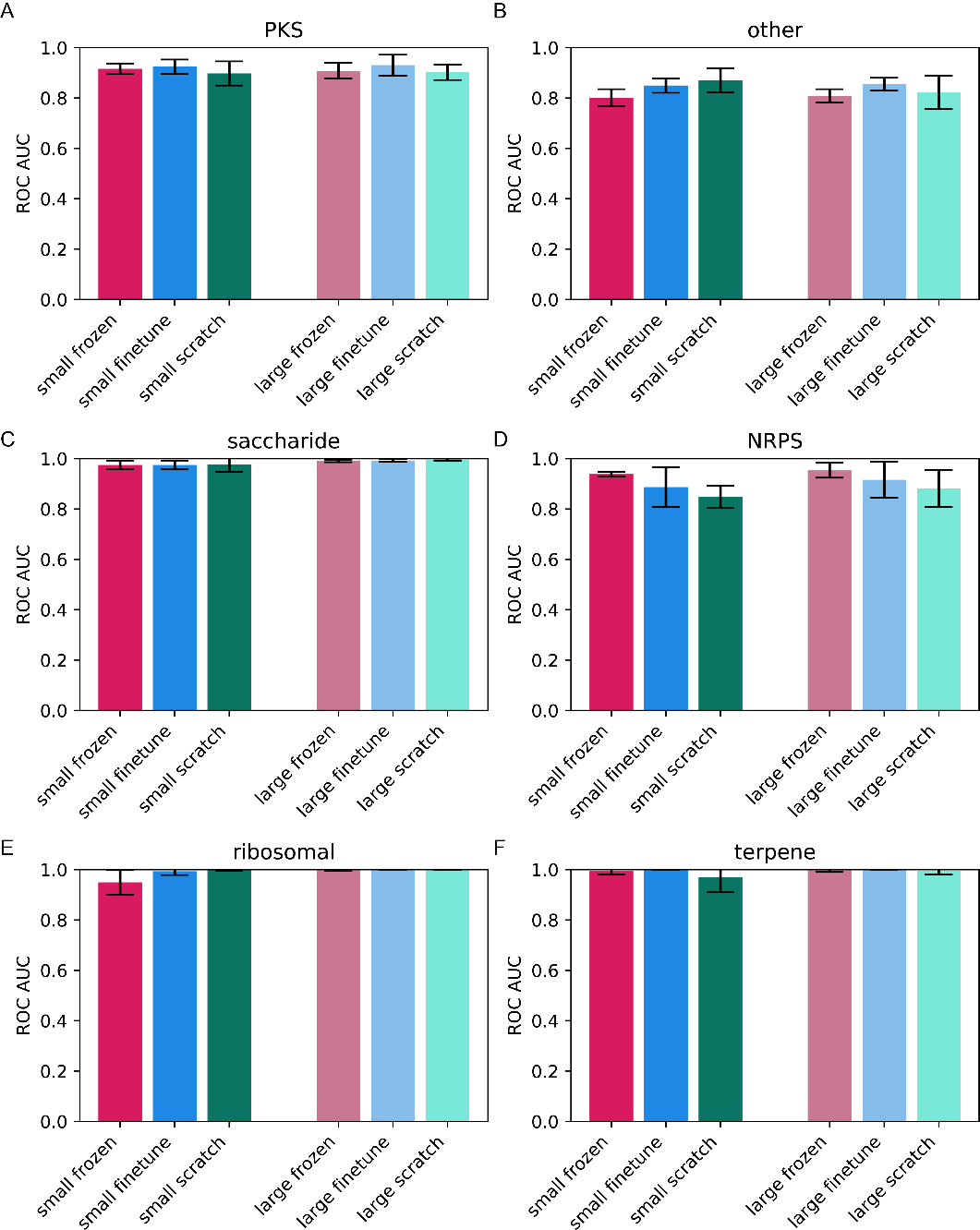

**Figure S3. AUROC for MIBiG classifications, test set with 20-40% similarity to pretraining set.** Classes that are not shown did not have enough BGCs in the 20-40% similarity range to calculate metrics. Error bars indicate standard deviation between models trained with different random seeds.

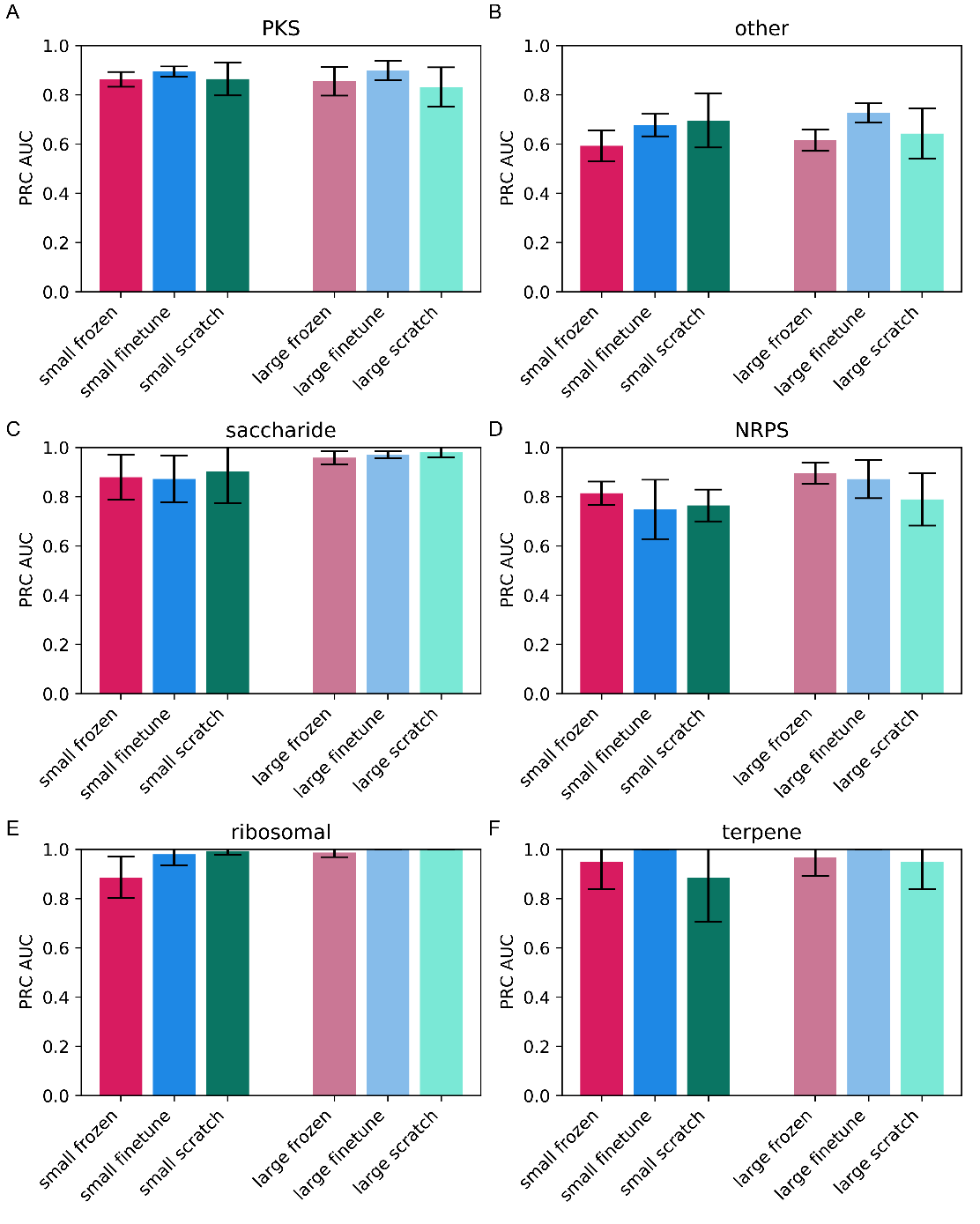

**Figure S4. AUPRC for MIBiG classifications, test set with 20-40% similarity to pretraining set.** Classes that are not shown did not have enough BGCs in the 20-40% similarity range to calculate metrics. Error bars indicate standard deviation between models trained with different random seeds.

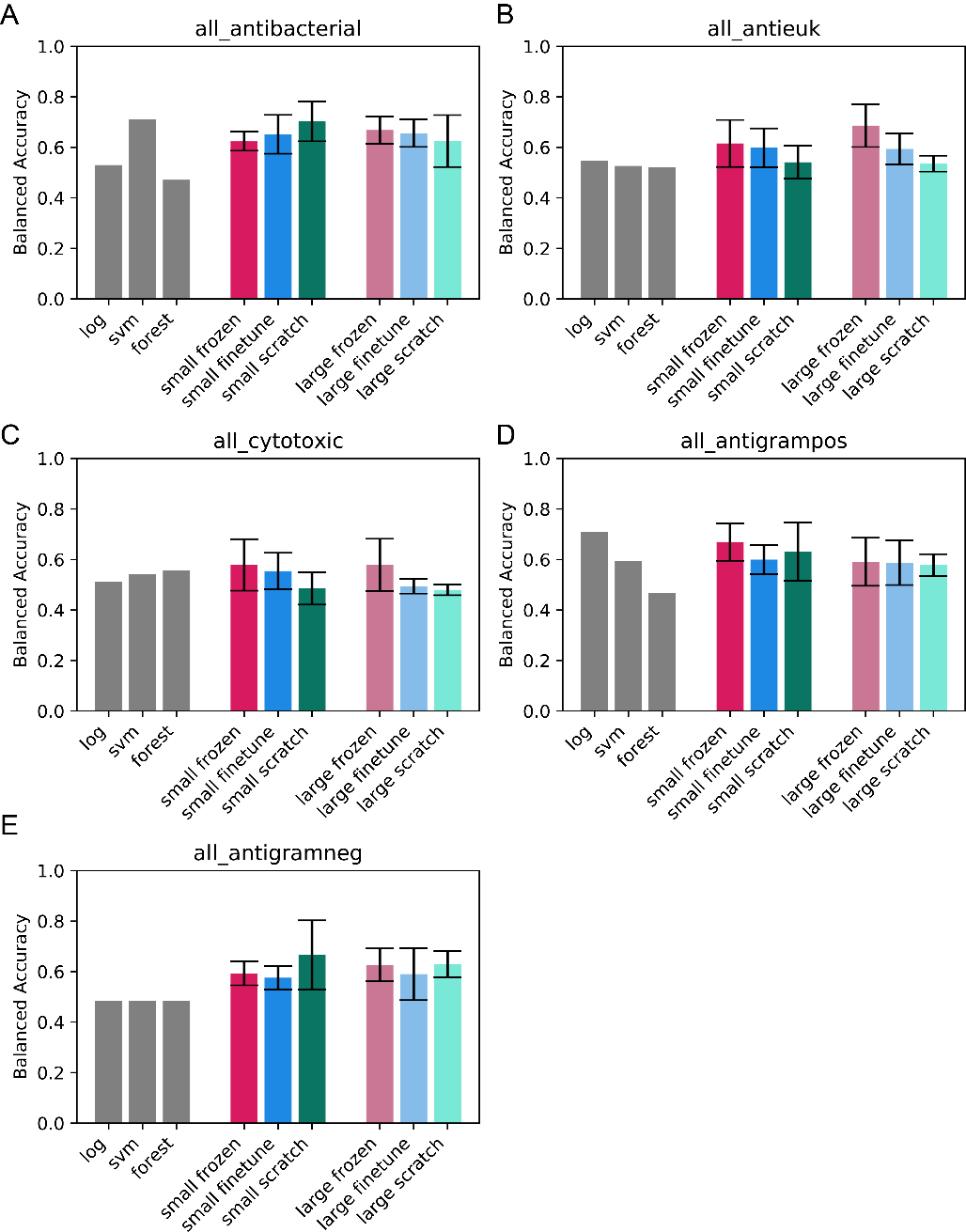

**Figure S5. Balanced accuracy for model trained on all activities applied to holdout set BGCs not identified by antiSMASH.** Grey bars indicate score from models from previous work, numbers reproduced from previous work.^1^ Error bars indicate standard deviation across five models.

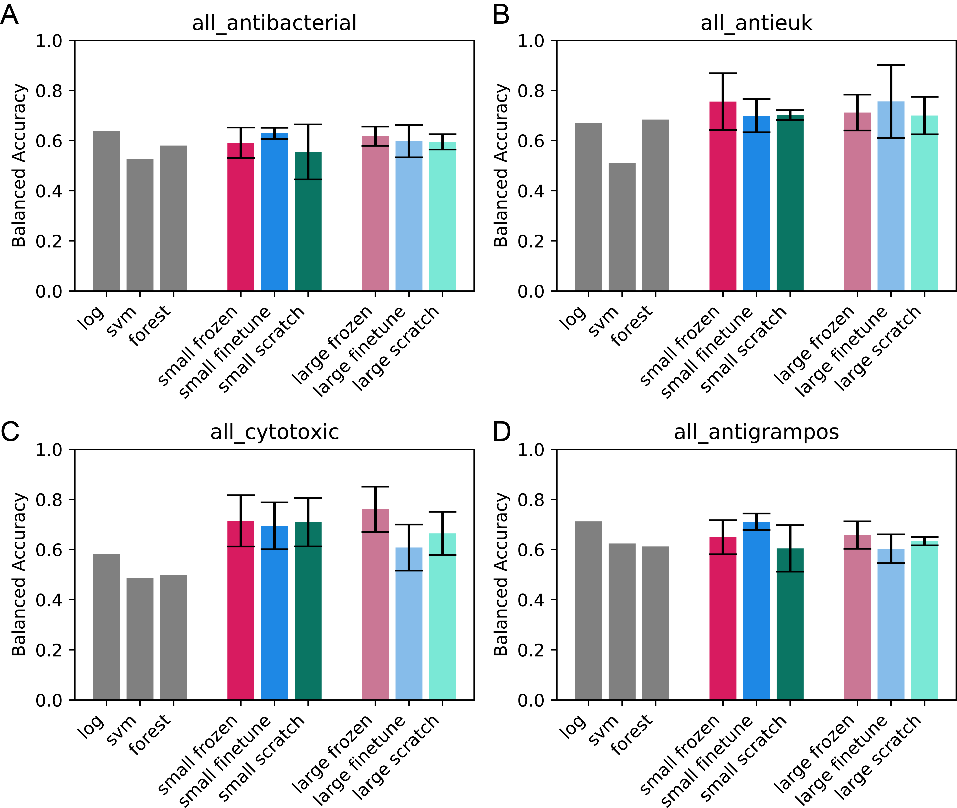

**Figure S6. Balanced accuracy for model trained on all activities applied to holdout set BGCs with no knownclusterblast hit.** Grey bars indicate score from models from previous work, numbers reproduced from previous work.^1^ Error bars indicate standard deviation across five models.

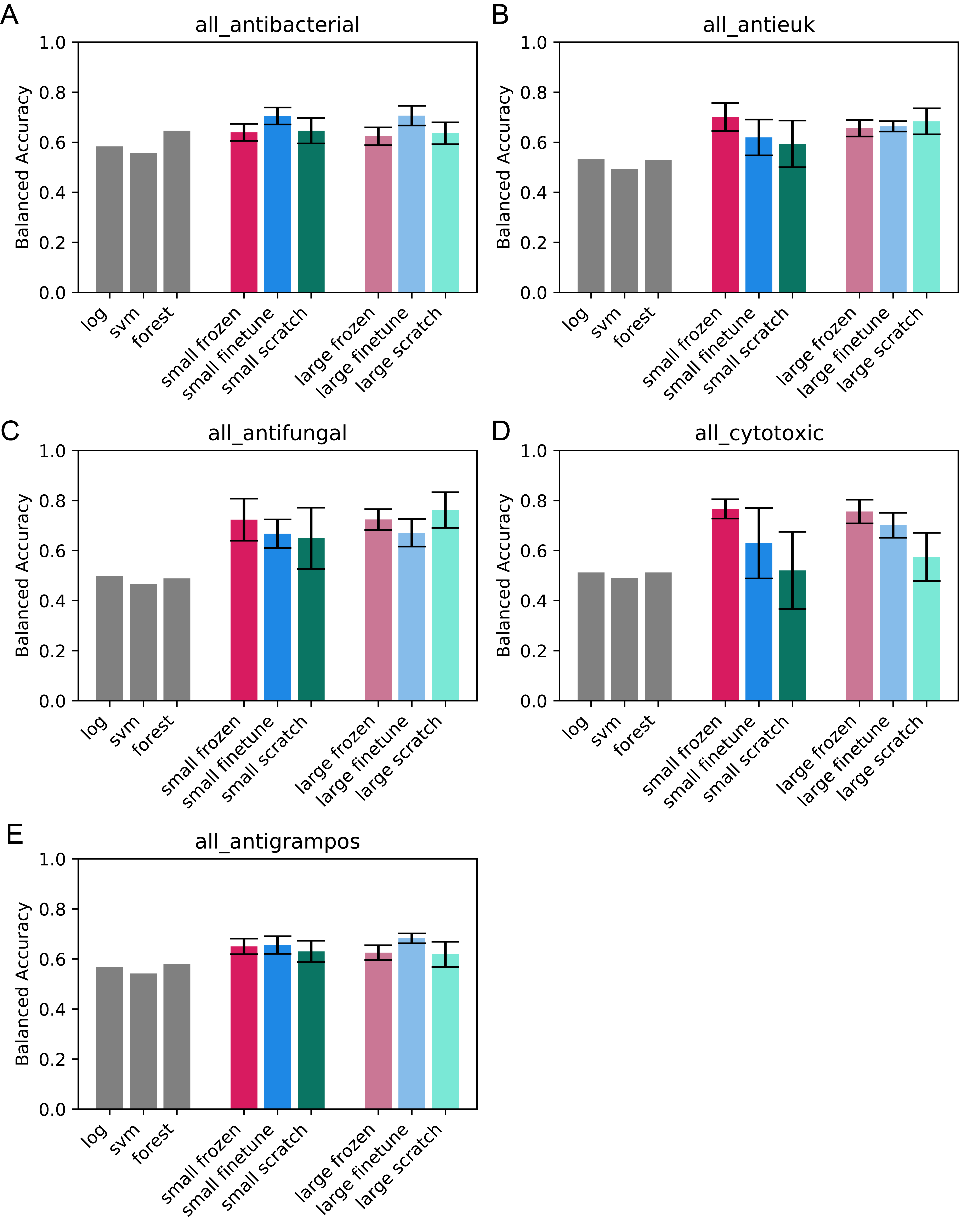

**Figure S7. Balanced accuracy for model trained on all activities applied to holdout set BGCs with knownclusterblast scores between 0 and 25% to training set BGCs.** Grey bars indicate score from models from previous work, numbers reproduced from previous work.^1^ Error bars indicate standard deviation across five models.

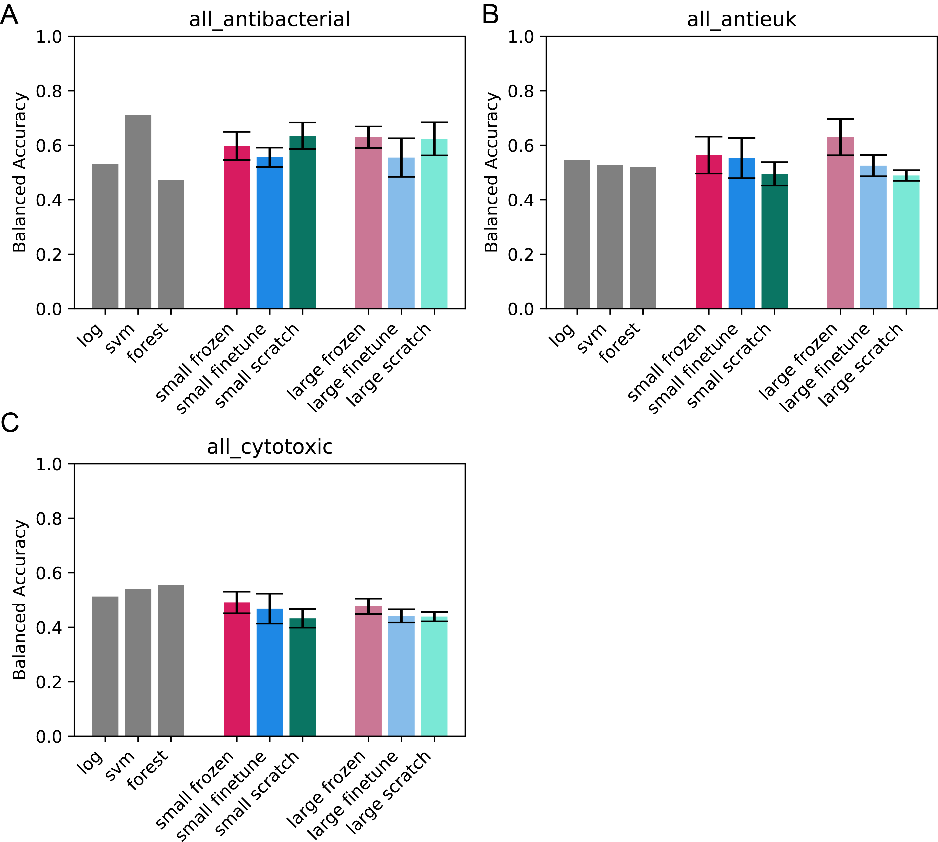

**Figure S8. Balanced accuracy for model trained on activities excluding type of bacterial target applied to holdout set BGCs not identified by antiSMASH.** Grey bars indicate score from models from previous work, numbers reproduced from previous work.^1^ Error bars indicate standard deviation across five models.

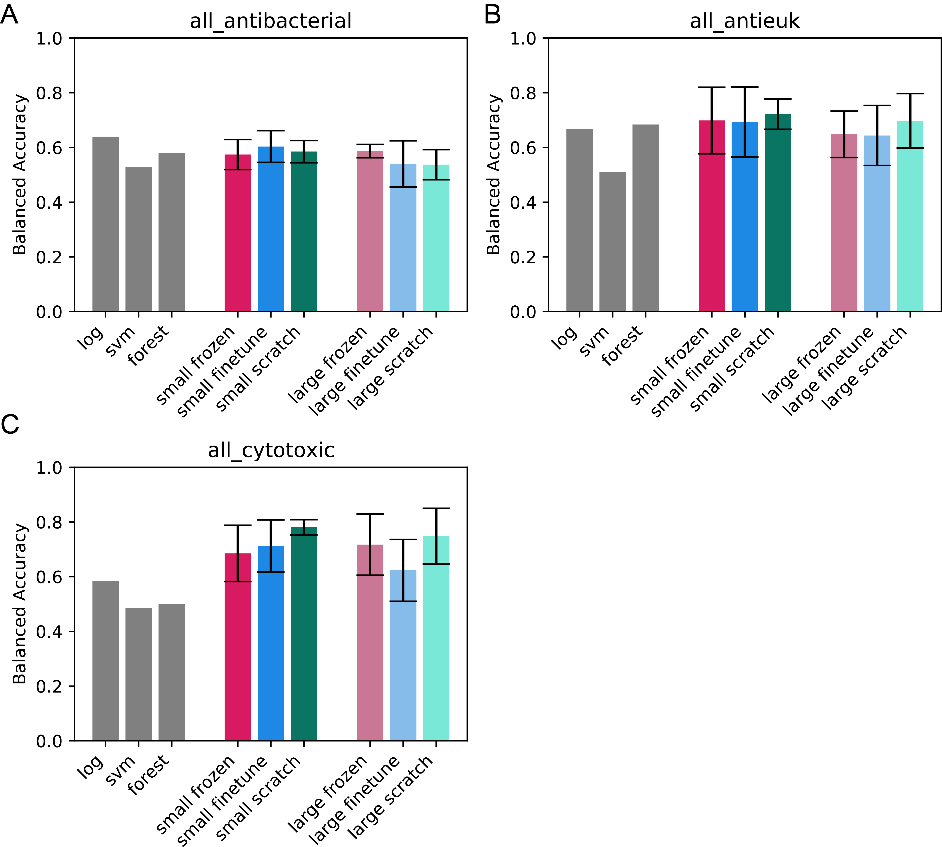

**Figure S9. Balanced accuracy for model trained on activities excluding type of bacterial target applied to holdout set BGCs with no knownclusterblast hit.** Grey bars indicate score from models from previous work, numbers reproduced from previous work.^1^ Error bars indicate standard deviation across five models.

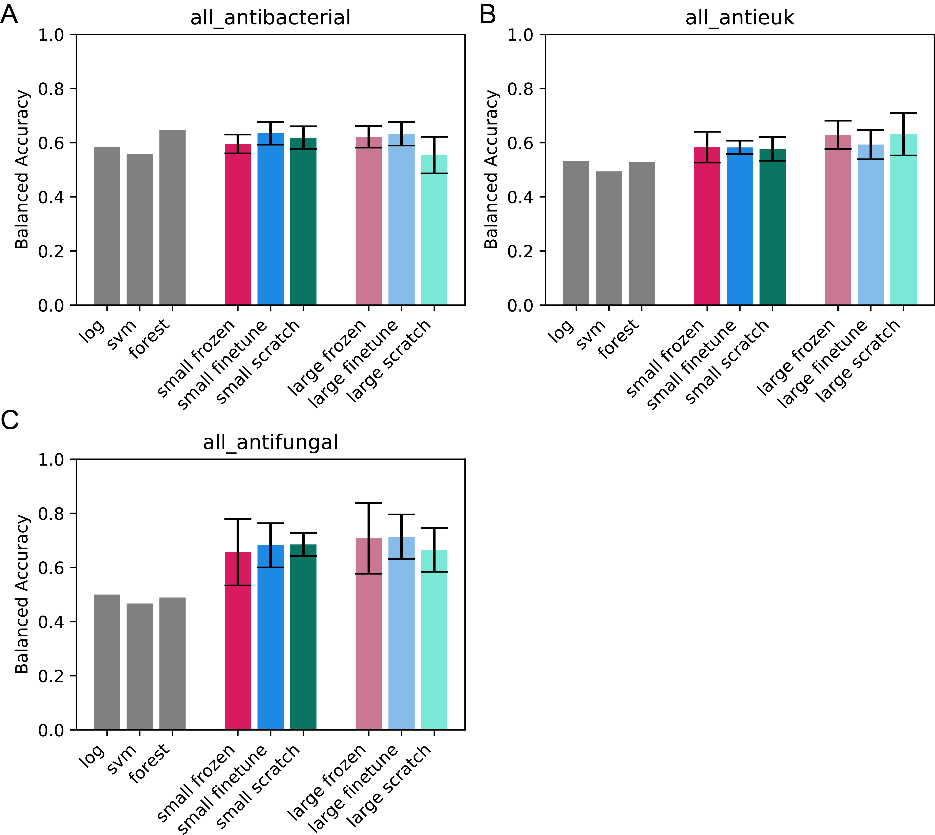

**Figure S10. Balanced accuracy for model trained on activities excluding type of bacterial target applied to holdout set BGCs with knownclusterblast scores between 0 and 25% to training set BGCs.** Grey bars indicate score from models from previous work, numbers reproduced from previous work.^1^ Error bars indicate standard deviation across five models.

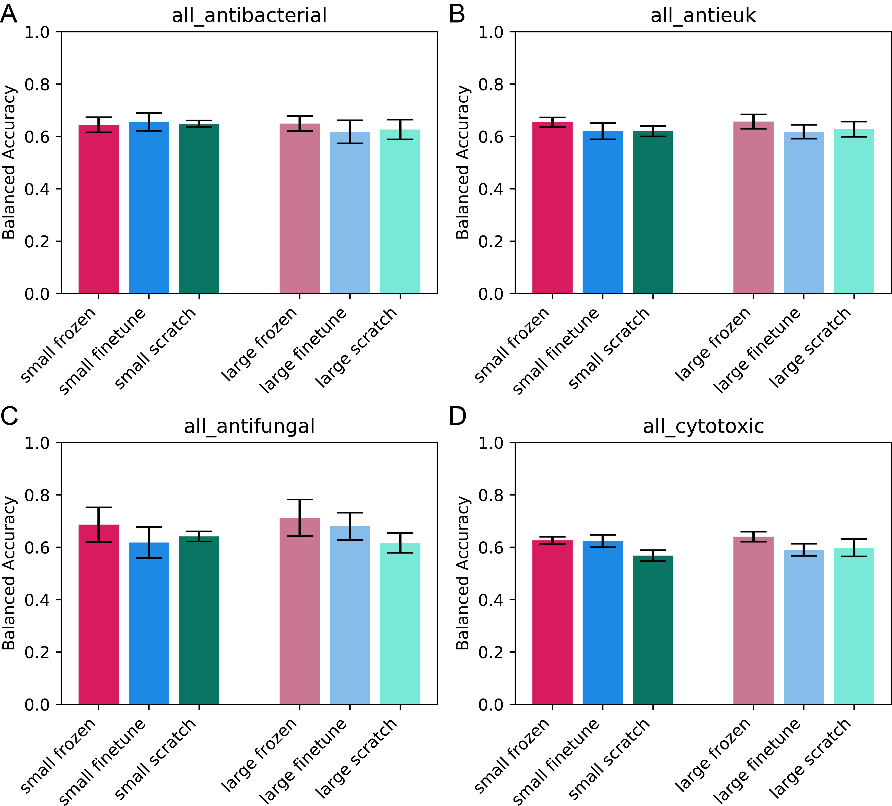

**Figure S11. Balanced accuracy for model trained on activities excluding type of bacterial target on entire hold outset.** Error bars indicate standard deviation across five models.

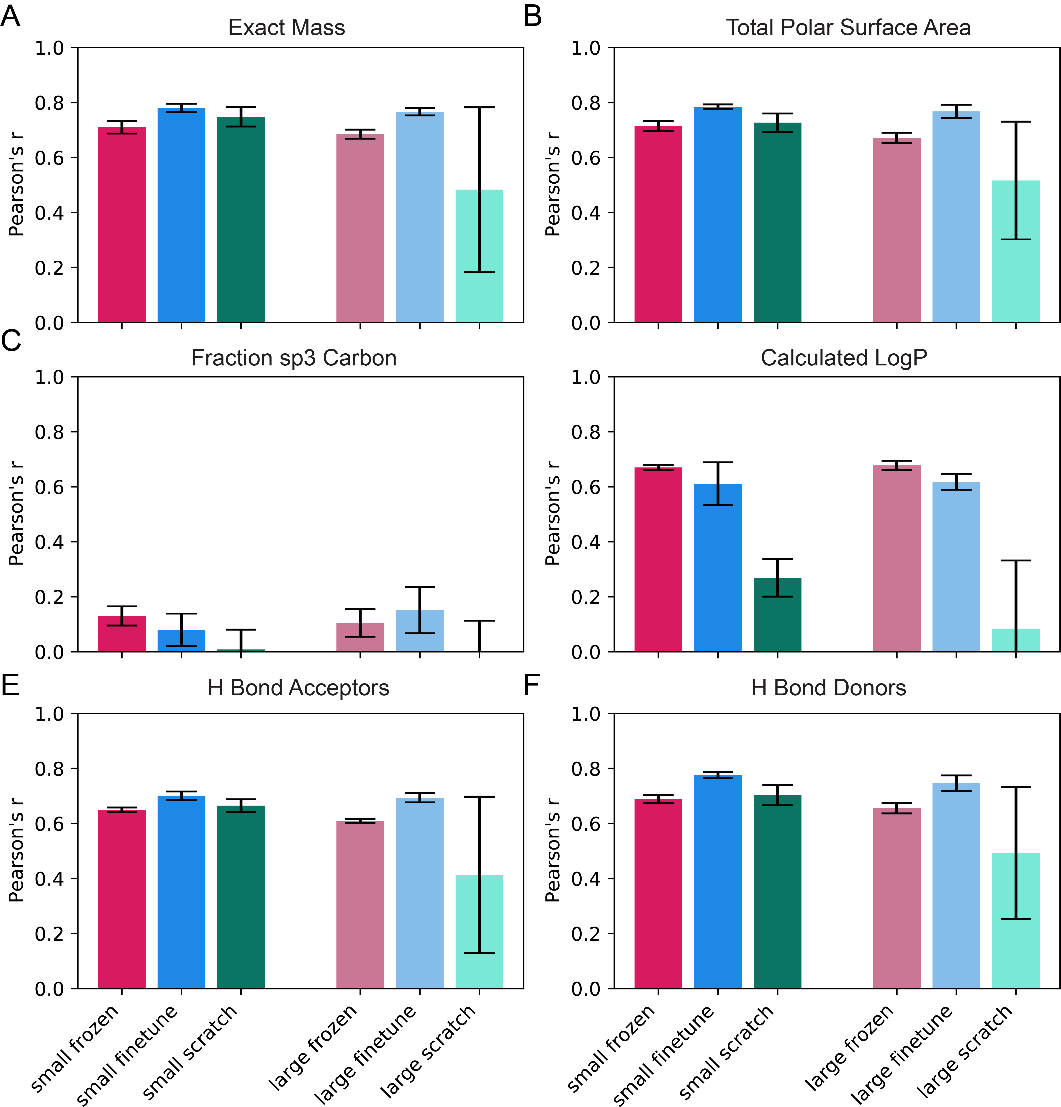

**Figure S12. Prediction of minimum descriptor values.** Error bars indicate standard deviation across five models.

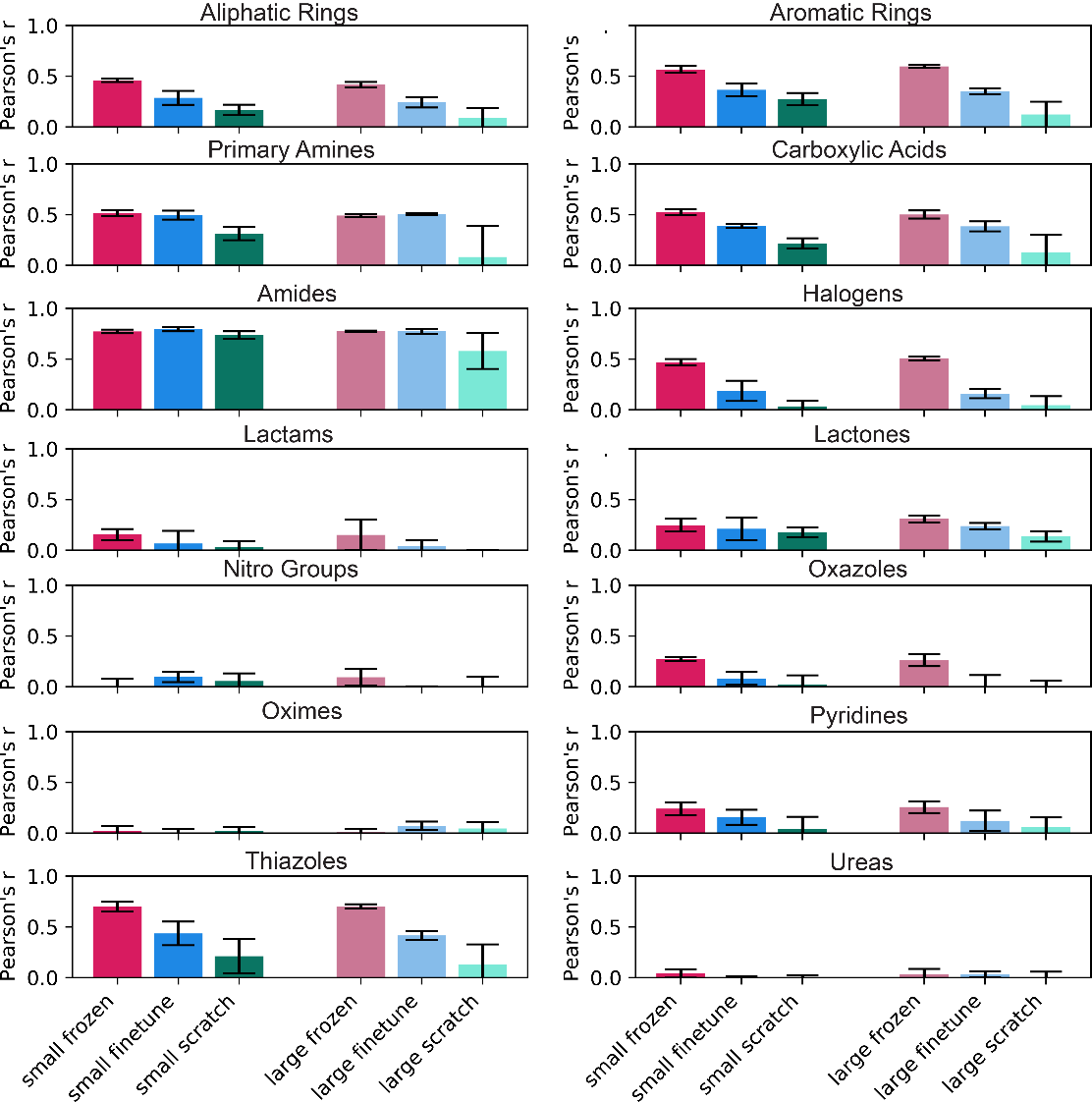

**Figure S13. Prediction of minimum number of functional group across products of a BGC.** Bars indicate Pearson’s correlation coefficient, error bars indicate standard deviation across five models.

**Table S5. Fingerprint Ranking Comparison, BGC to Molecule**

| Fingerprint method | Fingerprint size | Class weights | Average cosine rank | Average Tanimoto Rank |
| --- | --- | --- | --- | --- |
| Atom pair | 2048 | no | 310.2 | 187.2 |
| Atom pair | 4096 | no | 291.2 | 194.7 |
| Atom pair | 8192 | no | 276.9 | 190.0 |
| Atom pair | 2048 | yes | 844.5 | 845.0 |
| Atom pair | 4096 | yes | 843.6 | 843.9 |
| Atom pair | 8192 | yes | 843.5 | 843.9 |
| Functional morgan | 2048 | no | 223.1 | 256.3 |
| Functional morgan | 4096 | no | 239.7 | 272.6 |
| Functional morgan | 8192 | no | 272.6 | 286.4 |
| Functional morgan | 2048 | yes | 578.8 | 704.6 |
| Functional morgan | 4096 | yes | 536.9 | 673.4 |
| Functional morgan | 8192 | yes | 518.1 | 655.9 |
| Morgan | 2048 | no | **202.0** | **182.9** |
| Morgan | 4096 | no | 224.5 | 217.4 |
| Morgan | 8192 | no | 246.6 | 232.9 |
| Morgan | 2048 | yes | 710.0 | 799.4 |
| Morgan | 4096 | yes | 630.5 | 748.6 |
| Morgan | 8192 | yes | 572.8 | 708.1 |
| RdKit | 2048 | no | 351.5 | 290.2 |
| RdKit | 4096 | no | 268.0 | 217.5 |
| RdKit | 8192 | no | 254.8 | 243.2 |
| RdKit | 2048 | yes | 757.7 | 758.0 |
| RdKit | 4096 | yes | 759.4 | 759.6 |
| RdKit | 8192 | yes | 759.7 | 760.1 |
| Topological torsion | 2048 | no | 274.3 | 257.7 |
| Topological torsion | 4096 | no | 292.9 | 303.0 |
| Topological torsion | 8192 | no | 316.4 | 309.5 |
| Topological torsion | 2048 | yes | 734.6 | 798.7 |
| Topological torsion | 4096 | yes | 681.1 | 764.8 |
| Topological torsion | 8192 | yes | 647.2 | 741.0 |
| MACCS | NA | no | 217.3 | 203.8 |
| MACCS | NA | yes | 748.1 | 776.9 |

**Table S6. Fingerprint Ranking Comparison, Molecule to BGC**

| Fingerprint method | Fingerprint size | Class weights | Average cosine rank | Average Tanimoto Rank |
| --- | --- | --- | --- | --- |
| Atom pair | 2048 | no | 89.3 | 162.8 |
| Atom pair | 4096 | no | 82.0 | 155.0 |
| Atom pair | 8192 | no | 81.7 | 159.0 |
| Atom pair | 2048 | yes | 569.2 | 744.9 |
| Atom pair | 4096 | yes | 565.8 | 602.6 |
| Atom pair | 8192 | yes | 585.9 | 602.1 |
| Functional morgan | 2048 | no | 86.1 | 162.3 |
| Functional morgan | 4096 | no | 90.8 | 175.6 |
| Functional morgan | 8192 | no | 96.5 | 175.3 |
| Functional morgan | 2048 | yes | 328.7 | 511.8 |
| Functional morgan | 4096 | yes | 308.5 | 488.3 |
| Functional morgan | 8192 | yes | 301.3 | 470.7 |
| Morgan | 2048 | no | 69.7 | 143.4 |
| Morgan | 4096 | no | 74.1 | 149.6 |
| Morgan | 8192 | no | 80.6 | 150.0 |
| Morgan | 2048 | yes | 359.7 | 563.6 |
| Morgan | 4096 | yes | 302.0 | 522.6 |
| Morgan | 8192 | yes | 284.5 | 494.8 |
| RdKit | 2048 | no | 111.0 | 212.9 |
| RdKit | 4096 | no | 96.6 | 220.1 |
| RdKit | 8192 | no | 89.7 | 201.0 |
| RdKit | 2048 | yes | 559.5 | 997.9 |
| RdKit | 4096 | yes | 561.6 | 997.9 |
| RdKit | 8192 | yes | 498.1 | 998.0 |
| Topological torsion | 2048 | no | 80.5 | 230.3 |
| Topological torsion | 4096 | no | 82.0 | 255.4 |
| Topological torsion | 8192 | no | 95.2 | 237.1 |
| Topological torsion | 2048 | yes | 374.4 | 536.0 |
| Topological torsion | 4096 | yes | 320.5 | 503.1 |
| Topological torsion | 8192 | yes | 312.1 | 497.4 |
| MACCS | NA | no | **67.5** | **106.0** |
| MACCS | NA | yes | 560.2 | 626.3 |

**Table S7. Metrics for prediction of fingerprints.** Average metrics across five models trained with different random seeds for each model type, standard deviation in parentheses. Best model in bold. AUROC calculation is limited to classes with positive labels for the test set. AP indicates average precision.

|  | Micro Averaging | | | | Macro Averaging | | | |
| --- | --- | --- | --- | --- | --- | --- | --- | --- |
| Model | AP | AUROC | Recall | Precision | AP | AUROC | Recall | Precision |
|  | Morgan 2048 - Entire Test Set | | | | | | | |
| Small frozen | 0.23 (0.00) | 0.86 (0.00) | 0.29 (0.00) | **0.71 (0.01)** | 0.08 (0.00) | 0.74 (0.00) | 0.07 (0.00) | 0.15 (0.01) |
| Small finetune | 0.27 (0.02) | 0.85 (0.00) | 0.40 (0.02) | 0.62 (0.02) | 0.14 (0.01) | 0.75 (0.01) | 0.17 (0.02) | 0.27 (0.02) |
| Small scratch | 0.24 (0.00) | 0.85 (0.00) | 0.36 (0.01) | 0.61 (0.01) | 0.12 (0.00) | 0.74 (0.00) | 0.14 (0.01) | 0.23 (0.01) |
| Large frozen | 0.25 (0.00) | **0.87 (0.00)** | 0.33 (0.01) | 0.70 (0.01) | 0.09 (0.00) | **0.76 (0.00)** | 0.09 (0.00) | 0.19 (0.01) |
| Large finetune | **0.29 (0.01)** | 0.85 (0.00) | **0.43 (0.01)** | 0.64 (0.01) | **0.17 (0.01)** | 0.75 (0.00) | **0.20 (0.01)** | **0.31 (0.01)** |
| Large scratch | 0.24 (0.01) | 0.85 (0.00) | 0.38 (0.01) | 0.26 (0.02) | 0.13 (0.01) | 0.73 (0.01) | 0.16 (0.01) | 0.26 (0.02) |
|  | MACCS – Entire Test Set | | | | | | | |
| Small frozen | 0.71 (0.00) | 0.93 (0.00) | 0.79 (0.00) | 0.81 (0.00) | 0.44 (0.00) | 0.81 (0.00) | 0.47 (0.00) | 0.55 (0.01) |
| Small finetune | 0.73 (0.00) | 0.93 (0.00) | 0.80 (0.00) | **0.83 (0.00)** | 0.46 (0.01) | 0.81 (0.00) | 0.50 (0.00) | 0.57 (0.01) |
| Small scratch | 0.70 (0.01) | 0.92 (0.00) | 0.78 (0.00) | 0.81 (0.00) | 0.44 (0.01) | 0.78 (0.01) | 0.48 (0.01) | 0.54 (0.01) |
| Large frozen | 0.71 (0.00) | 0.93 (0.00) | 0.78 (0.01) | 0.82 (0.01) | 0.44 (0.00) | 0.81 (0.00) | 0.46 (0.01) | 0.56 (0.01) |
| Large finetune | **0.73 (0.00)** | **0.93 (0.00)** | **0.81 (0.01)** | 0.83 (0.00) | **0.47 (0.00)** | **0.82 (0.01)** | **0.52 (0.00)** | **0.59 (0.01)** |
| Large scratch | 0.70 (0.01) | 0.92 (0.00) | 0.78 (0.00) | 0.81 (0.01) | 0.44 (0.01) | 0.79 (0.02) | 0.48 (0.01) | 0.55 (0.01) |
